# Kymograph classifiers predict CMT2A disease-relevant mitochondrial motility phenotypes in iPSC motor neurons

**DOI:** 10.64898/2026.03.16.712192

**Authors:** Adam C Weiner, Bria L Macklin, Leo Epstein, Kaitlin R Kelly, Bruce R Conklin, Barbara E Engelhardt

## Abstract

Charcot-Marie-Tooth disease type 2A (CMT2A) is an autosomal dominant peripheral neuropathy caused by *MFN2* mutations that impair mitochondrial trafficking, leading to progressive motor neuron degeneration and loss of motor function in patients. To generate a defined disease model, we engineered one of the most severe mutations, *MFN*2^R364W^, into a healthy wild-type iPSC background, and then we differentiated isogenic mutant and healthy lines into motor neurons. Historically, mitochondrial trafficking phenotypes have been quantified using discrete, manually annotated metrics, an approach that is difficult to scale. To address this, we developed an image classifier that predicts disease phenotype from kymographs, an image representation of mitochondrial movement along axons automatically derived from videos. Our method, KymoClass, discriminates healthy *MFN*2^WT^ from diseased *MFN*2^R364W^ iPSC motor neurons more accurately than summary statistics extracted from kymographs, such as mitochondrial speed and stationary fraction, and remains accurate on a held-out biological replicate. Interpretation of classifier attention scores and vision transformer patch embeddings reveals a previously undescribed subtype of stationary mitochondria, which we call “jitter”, showing small-amplitude back-and-forth motion, which associates most strongly with R364W kymographs. Applied to three *MFN2* mutations excluded from training—T105A, W740S, and R94W—the classifier assigns pathogenicity scores consistent with reported clinical severity. This approach may be used to enable high-throughput screens in which axonal organelle trafficking drives disease pathogenesis.

## Introduction

Charcot-Marie-Tooth (CMT) disease is among the most commonly inherited neurological disorders, with an estimated population prevalence of 1:2500 to 1:10000^1^. CMT type 2A (CMT2A), one of the most frequent subtypes of CMT, can be caused by more than 100 different missense mutations in the *MFN2* gene that each exhibit different degrees of pathogenicity^2^. CMT2A is primarily classified as an autosomal-dominant inherited neuropathy, meaning that patients most commonly have one wild-type (WT) allele and one mutant allele at the *MFN2* gene^3–5^. *MFN2* encodes a ubiquitous GTPase that regulates the fusion of the outer mitochondrial membrane, which is essential for proper mitochondrial trafficking. Severe *MFN2* mutations lead to an early-onset disease with rapid degeneration of the peripheral motor and sensory nervous systems, whereas some mutations result in a milder clinical expression. There is an ongoing gap in our understanding of how different mutations affect mitochondria and the resulting cellular phenotypes^6^.

Mitochondrial trafficking is integral to cellular health, allowing cells to respond to energetic demands and changing nutrient environments. Neurons face unique energetic needs, requiring high concentrations of mitochondria at the synapse site, while also needing to maintain a specific distribution of mitochondria throughout lengthy axons^7,8^. Since mutations in *MFN2* were first linked to CMT2A in 2004^3^, several groups have studied the genotype-to-phenotype correlations across different mutations. Using non-neuronal cell types as model systems, early studies found differences in mitochondrial network formation in *MFN*2^mut^ human cells^9–12^; however, conflicting results have been reported, potentially due to *MFN1* expression in these cell types^13^ and differing mitochondrial metabolic activity when mutant *MFN2* is overexpressed^14^. This work was supported by mouse studies, which found abnormal mitochondrial trafficking patterns in several different mutations^15–17^. The differences across studies were correlated with the protein domain of the mutated amino acid residue^18–20^, suggesting that different types of mitochondrial dysfunction can drive CMT2A pathology.

While these works provided insight into MFN2 biology, the advent of induced pluripotent stem cell (iPSC) technology has enabled researchers to generate high-quality model systems for CMT2A, deriving phenotypic motor neurons from patient or mutation-engineered iPSCs^21–24^. Using iPSC-derived motor neurons, researchers have reported a reduction in mitochondrial trafficking along axons in *MFN*2^mut^ neurons as compared to *MFN*2^WT^ neurons^22–24^. Cell models derived from patient iPSCs are extremely valuable for studying disease phenotypes; however, a limitation of these studies is the lack of isogenic WT controls. This is a critical gap, as the genetic background of iPSCs has been shown to affect pluripotency and differentiation potential^25–27^, which can confound downstream phenotyping of iPSC derivatives. An additional area of opportunity is the generation of a platform that studies multiple *MFN2* mutations to develop a clearer picture of mitochondrial dysfunction in CMT2A. Here, we introduce an isogenic panel of four different *MFN2* mutants, selected by range in clinical severity along a single axis and located in different functional domains of the protein, engineered into a fully characterized iPSC WT background, such that each cell line differs only at the site of the mutation.

Time-lapse fluorescence imaging is often performed on *in vitro* samples of CMT2A motor neurons to quantify mitochondrial trafficking along axons. In these experiments, kymographs are often used to measure mitochondrial fluorescence intensity across each axon pixel and time frame (Extended Data Fig. 1). While kymographs have been used for many years to represent mitochondrial dynamics as two-dimensional images, most researchers analyzed kymographs using manual tracing of individual mitochondrial tracks^28,29^. Recent methods, such as KymographClear^30^, KymoButler^31^, and TrackMate-Kymograph^32^, perform automated segmentation of kymograph tracks. These methods enable researchers to compute easily interpretable statistics such as mitochondrial speed, pausing events, and total displacement^23,33^; however, they currently do not estimate line thickness (mitochondrial aggregate size), and segmentation errors can confound the estimation of summary statistics.

In this work, we draw from recent advancements in computer vision^34,35^ to develop KymoClass: an interpretable image classifier for scoring the phenotype of kymographs obtained from fluorescence time-lapse imaging of iPSC motor neurons. Using *MFN*2^WT^ and *MFN*2^R364W^ iPSC motor neurons, our classifier predicts *MFN2* genotype from kymograph images on both internal validation and held-out test data, effectively scoring the disease severity of each well based on mitochondrial movement. We then analyze KymoClass’s attention scores and vision transformer (ViT) patch embeddings to interpret which mitochondrial trafficking properties are associated with genotype. Finally, we show that KymoClass generalizes to *MFN2* genotypes that it never saw during training, scoring the pathogenicity of the T105A, W740S, and R94W mutations in a manner consistent with their reported clinical severity. To our knowledge, this work is the first to train image classifiers on kymograph summaries of video data to predict, score, and interpret disease-relevant cellular phenotypes. This study is also the first to directly compare mitochondrial motion across multiple *MFN2* mutants and present a model for scoring *MFN2* mutations that correlate with clinical phenotype. The resulting KymoClass method enables fast and precise analysis of large-scale screens for therapeutic approaches for CMT2A and other mitochondrial diseases.

## Results

### Generation of an isogenic series of *MFN2* mutant iPSCs

To generate a reproducible model of CMT2A pathology, we selected four *MFN2* missense mutations, R94W, R364W, T105A, and W740S for our iPSC model system. Each of these mutations falls within a different domain of the predicted MFN2 protein structure and correlates with known clinical phenotypes^36^. R364W and R94W have a severe, earlier age of onset of disease, and T105A and W740S mutants have a less profound effect on motor function and an older age of onset^6,37,38^ (Table 1; Supplementary Video 1). We nucleofected gRNA-HiFi Cas9^39^ ribonucleoprotein (RNP) and template oligos (Fig. 1a and Extended Data Fig. 2a) into a well-characterized WT iPSC control line (WTD)^40^. Single-cell clones were isolated and genotyped via NGS to confirm heterozygous insertion at the target site and clonality (Extended Data Fig. 2b).

**Figure 1.**
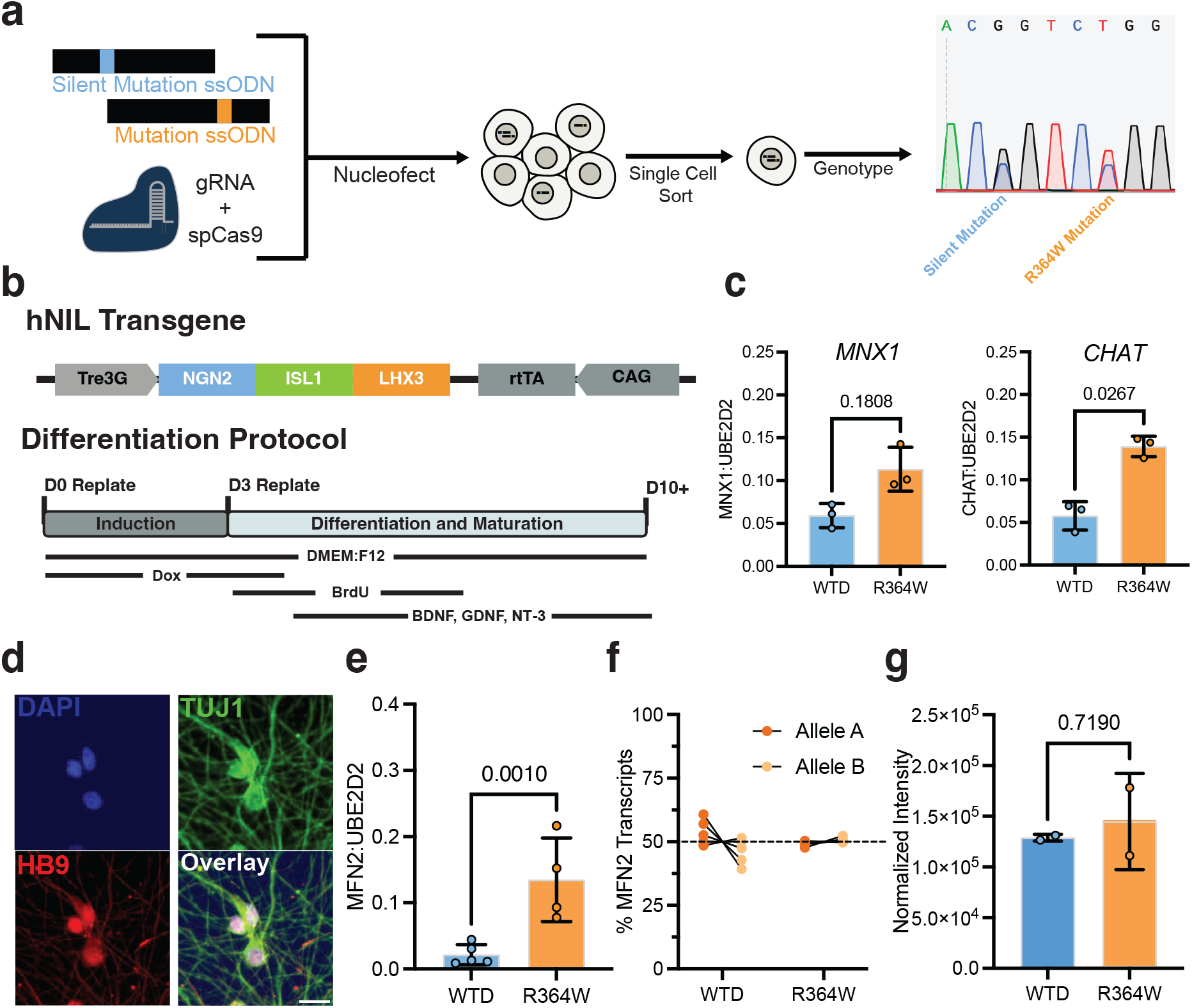
Generation of an isogenic series of *MFN*2^mut^ iPSCs. (A) Schematic outlining cell line engineering workflow. (B) i^3^ motor neuron transgene cassette and differentiation timeline. All assays were performed on day 10 of differentiation with WT and *MFN*2^R364W^ derivatives. (C) Digital PCR gene expression assay for motor neuron-specific transcription factor, *MNX1*, and the mature motor neuron marker, *CHAT*, normalized to *UBE2D2* expression. (D) Immunofluorescent-stained i^3^ motor neurons differentiated from the *MFN*2^R364W^ iPSC line. Scale bar = 20 *µ*m. (E) Digital PCR gene expression assay for total *MFN2* normalized to *UBE2D2* expression. (F) Digital PCR assay for relative expression of *MFN2* transcripts. (G) Total MFN2 protein expression measured by western blot, normalized to *β* -actin. For gene expression assays and western blot quantification (C, E, F, and G), a one-way ANOVA was performed, followed by Dunnett’s multiple comparison test against WT (all cell lines are shown in Extended Data Figs. 3 and 4).

**Table 1.**
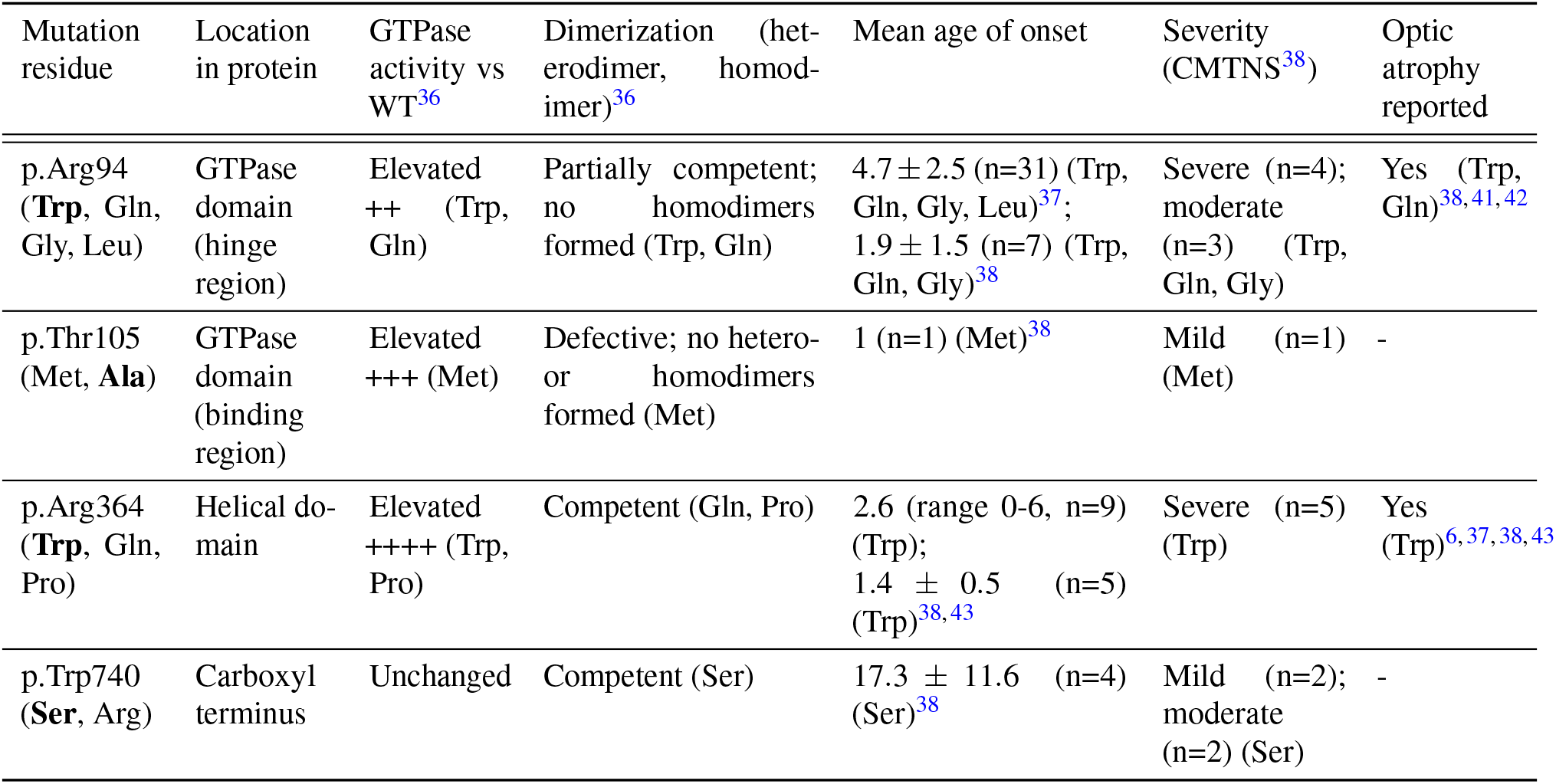
Summary of the *MFN2* missense mutations analyzed in this study. Average age of onset is listed in years. The CMT neuropathy score reported for T105A is that of the T105M variant at the same residue.

| Mutation residue | Location in protein | GTPase activity vs WT<sup>36</sup> | Dimerization (het-erodimer, homod-imer)<sup>36</sup> | Mean age of onset | Severity (CMTNS<sup>38</sup>) | Optic atrophy reported |
|---|---|---|---|---|---|---|
| p.Arg94 (**Trp**, Gln, Gly, Leu) | GTPase domain (hinge region) | Elevated ++ (Trp, Gln) | Partially competent; no homodimers formed (Trp, Gln) | 4.7 ± 2.5 (n=31) (Trp, Gln, Gly, Leu)<sup>37</sup>;<br>1.9 ± 1.5 (n=7) (Trp, Gln, Gly)<sup>38</sup> | Severe (n=4); moderate (n=3) (Trp, Gln, Gly) | Yes (Trp, Gln)<sup>38,41,42</sup> |
| p.Thr105 (Met, **Ala**) | GTPase domain (binding region) | Elevated +++ (Met) | Defective; no hetero- or homodimers formed (Met) | 1 (n=1) (Met)<sup>38</sup> | Mild (n=1) (Met) | - |
| p.Arg364 (**Trp**, Gln, Pro) | Helical do-main | Elevated ++++ (Trp, Pro) | Competent (Gln, Pro) | 2.6 (range 0-6, n=9) (Trp);<br>1.4 ± 0.5 (n=5) (Trp)<sup>38,43</sup> | Severe (n=5) (Trp) | Yes (Trp)<sup>6,37,38,43</sup> |
| p.Trp740 (**Ser**, Arg) | Carboxyl terminus | Unchanged | Competent (Ser) | 17.3 ± 11.6 (n=4) (Ser)<sup>38</sup> | Mild (n=2); moderate (n=2) (Ser) | - |

The WTD parental line used carries a previously integrated inducible cassette encoding *NGN2*, *ISL1*, and *LHX3* in the *CLYBL* locus, enabling rapid differentiation into motor neurons (Fig. 1b). Using the i^3^ motor neuron differentiation protocol, *MFN*2^mut^ clones were differentiated to day 10, then characterized for the motor neuron markers *MNX1* and *CHAT* by gene-expression digital PCR (Fig. 1c and Extended Data Fig. 3a). Nuclear localization of HB9, encoded by *MNX1*, was confirmed via immunohistochemistry with a beta-tubulin co-stain (Fig. 1d and Extended Data Fig. 3b). Measurement of total *MFN2* transcripts revealed an increased number of total transcripts in the *MFN*2^R364W^ clone (Fig. 1e and Extended Data Fig. 4a), consistent with previous reports^44^, with allelic expression balanced between the mutant and WT alleles (Fig. 1f and Extended Data Fig. 4b). However, this increase in *MFN2* mRNA did not correspond to a measurable change in protein level in the *MFN*2^R364W^ or any other *MFN*2^mut^ clone (Fig. 1g and Extended Data Fig. 4c).

### Encoding time-lapse motor neuron images as kymographs

To establish the platform, we began by analyzing the *MFN*2^R364W^ mutant, as this mutation is among the most severe and has been widely studied previously. Mutant and WT iPSCs were transduced with two lentiviral constructs: a mitochondrial localized dsRed2 and tubulin localized EGFP. For both cell lines, labeled cells were mixed 1:10 (labeled: unlabeled) with *MFN*2^R364W^ iPSCs with no label then differentiated to motor neurons (Fig. 2a). To image, each well from each biological replicate was subjected to 3 minutes of time-lapse fluorescence imaging, with each frame being 2 seconds apart (Fig. 2b). Through segmentation and filtering of the green channel, we obtained a mask of all axons present in each well (Fig. 2c). When computing the axon masks, we split axons into different segments at each axon–axon intersection to ensure that none of the detected segments contain multiple disconnected neurons. Finally, we computed mitochondrial fluorescence (red channel intensity) along each axon segment over time, creating kymographs where the x-axis represents the position along the axon and the y-axis represents time (Fig. 2d). Transforming these time-lapse videos into kymographs enabled us to distill mitochondrial motility dynamics into a single 2D image without having to control for well-level batch effects, such as axon seeding density, or axon-level batch effects, such as orientation.

**Figure 2.**
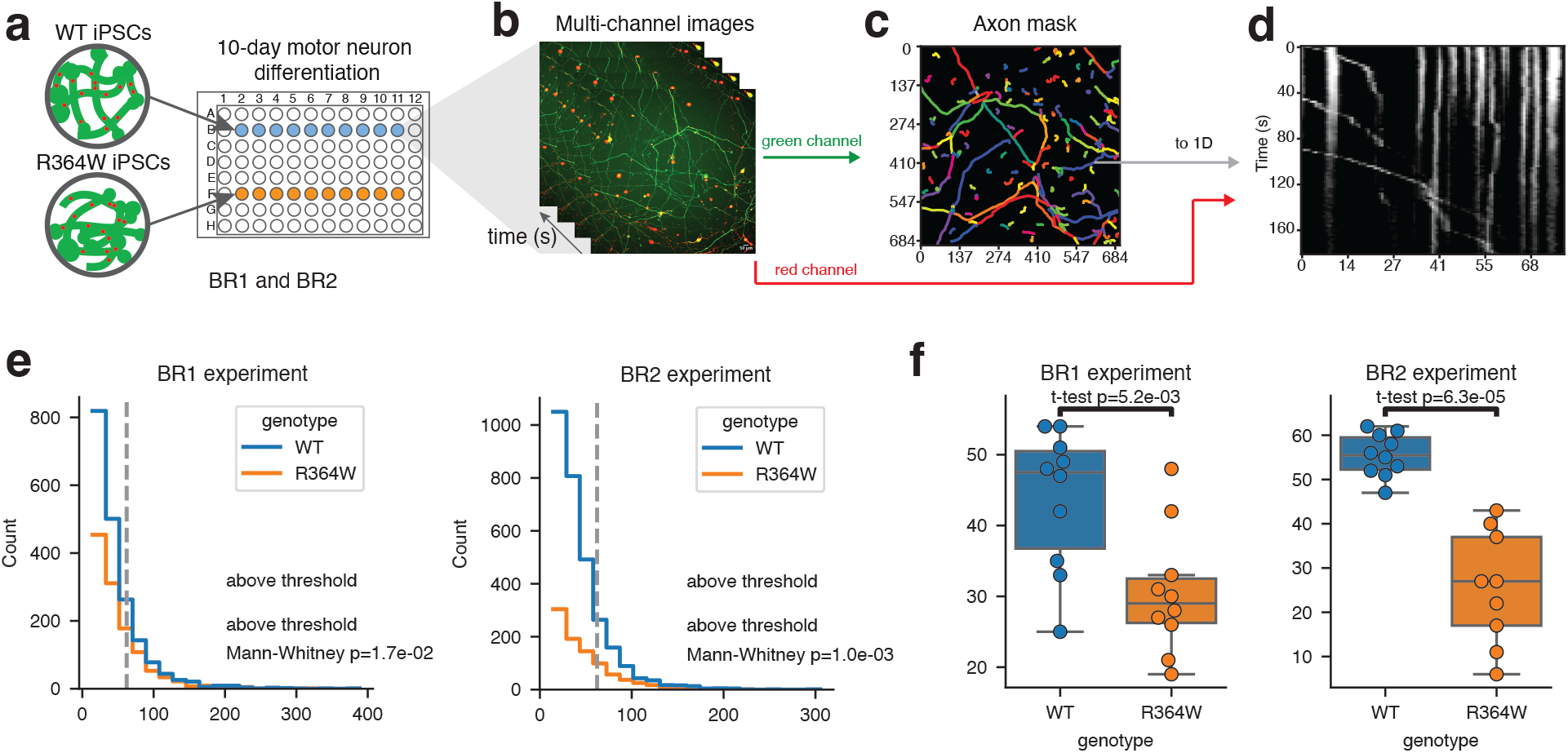
Using kymographs to represent mitochondrial motion along axons. (A-D) Experimental overview. (A) WT and R364W-mutant iPSCs were loaded onto two 96-well plates (one per biological replicate) and subjected to a 10-day doxycycline-inducible motor neuron differentiation protocol. (B) Time-lapse fluorescence imaging was performed at differentiation day 10. Each well was imaged every 2 seconds for a total of 3 minutes, capturing axons in the green channel and mitochondria in the red channel. (C) All green channel images of a given well were used to create an axon mask. (D) For each axon segment *α*, we produced a kymograph where white pixels correspond to high mitochondrial intensity and black pixels correspond to low mitochondrial intensity. (E) Distribution of axon segment lengths per genotype across the two biological replicates. Only axon segments *>* 62 *µ*m are included in downstream analysis. (F) Boxplot of the number of axon segments per genotype and biological replicate included in downstream analysis.

Before analyzing mitochondrial motility dynamics in our kymographs, we performed quality control filtering to ensure that all kymographs were generated from high-quality axon segments that contained mitochondria. First, we examined axon segment lengths across both biological replicate experiments, finding shorter axons in WT wells than in R364W wells across both experiments (Mann-Whitney tests *p ≤*1.7 *×* 10*^—^*^2^ in BR1, *p ≤*1.0 *×* 10*^—^*^3^ in BR2; Fig. 2e). Since shorter axon segments contain less information about mitochondrial motion and are more likely to be segmentation artifacts, we only considered kymographs coming from axons *>* 62 *µ*m in length for downstream analysis. After kymograph generation and filtering steps, we were left with a total of *n* = 1562 kymographs to analyze across all experiments. Moreover, there were more WT kymographs than R364W kymographs in total (two-sided t-tests *p ≤*5.2 *×* 10*^—^*^3^ in BR1, *p ≤*6.3 *×* 10*^—^*^5^ in BR2; Fig. 2f), likely due to higher viability of WT motor neurons compared to those with pathogenic mutations.

### Previous mitochondrial tracking software does not robustly differentiate *MFN2* genotypes

Our first approach for analyzing mitochondrial motility dynamics centered on the automated segmentation of individual mitochondrial tracks within each kymograph, which yields interpretable phenotypes such as velocity, acceleration, and pausing. We used KymoButler^31^ to segment all kymographs across the two experiments (Fig. 3a). Using a 0.1 *µ*m/s cutoff to stratify moving versus stationary mitochondria, consistent with thresholding strategies of previous publications^15^ (see Methods), KymoButler found no statistically significant difference in motility percentage between R364W and WT wells across both biological replicates (two-sided t-tests *p ≤*0.42 in BR1, *p ≤*0.10 in BR2; Fig. 3b). A logistic regression classifier trained on the BR1 motility percentages then predicted the genotype of the held-out BR2 wells poorly, indicating that this statistic does not behave as a genotype-specific phenotype in these samples (F1 = 0.67; Fig. 3c).

**Figure 3.**
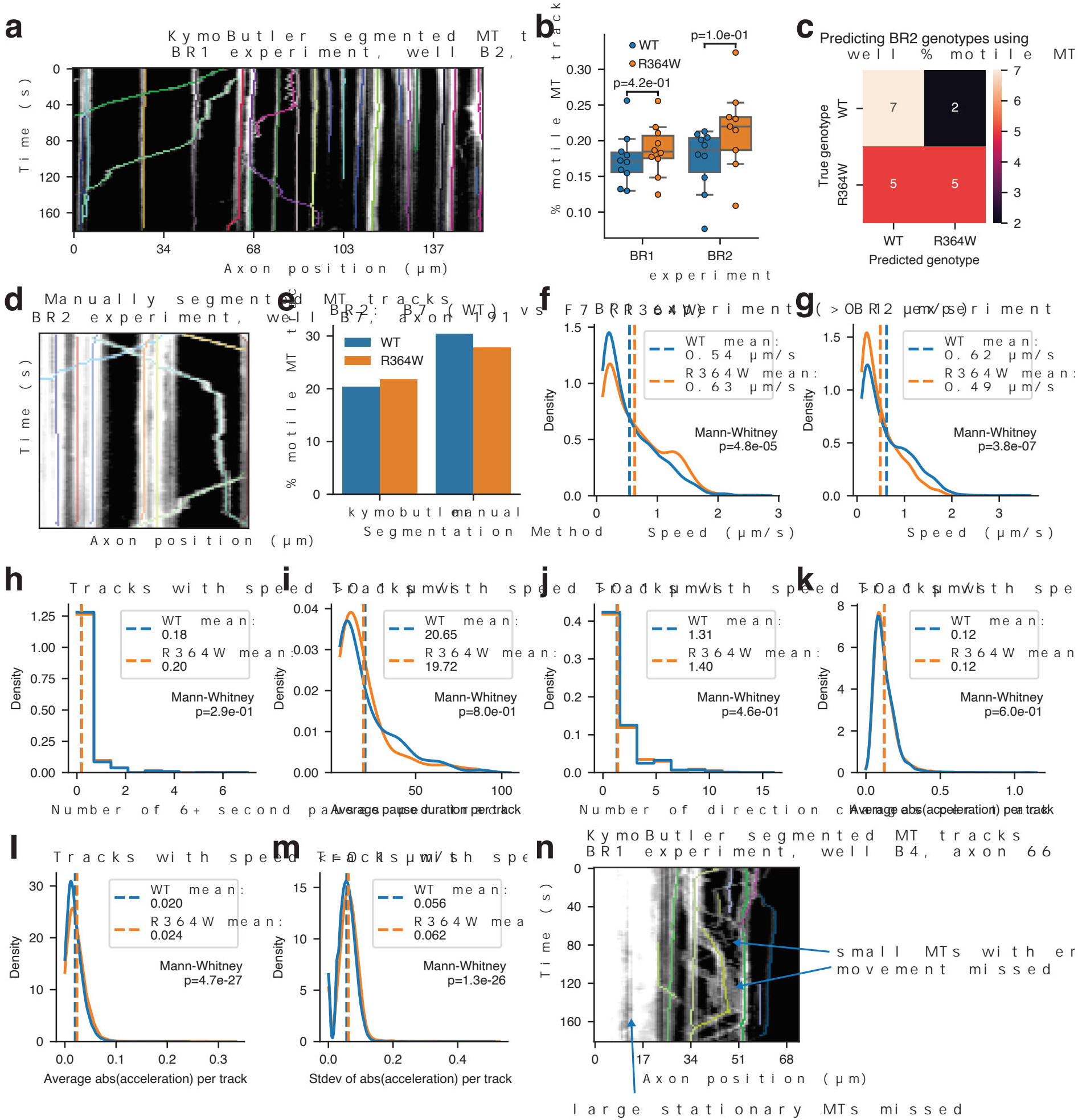
KymoButler track segmentation of kymographs. (A) KymoButler mitochondrial tracks superimposed on the raw kymograph for the BR1 experiment, well B02, axon 0. Each track is shown in a unique color. (B) Boxplot of the percent of motile mitochondria tracks within each well, split by genotype and biological replicate. A threshold of 0.1 *µ*m/s mean velocity is used to differentiate moving versus stationary tracks. (C) Confusion matrix for predicting the genotype of BR2 wells using a logistic regression classifier trained on the percent motile values from BR1 wells. (D) Manually segmented mitochondrial tracks superimposed on the raw kymograph for the BR2 experiment, well B07, axon 191. (E) Comparison of the percent motile mitochondria statistic between KymoButler and manual track segmentation approaches for wells B07 and F07 in the BR2 experiment. (F-G) Distribution of motile mitochondria track speeds split by genotype for the (F) BR1 experiment and (G) BR2 experiment. Mean speeds across all tracks are denoted by dashed vertical lines with color corresponding to genotype. (H-K) Distribution of motile mitochondria summary statistics across both experiments. (H) Number of pauses per track. Each pause must last at least 6 seconds in duration. (I) Average duration of each detected pause per track, excluding tracks with no 6+ second pauses. (J) Number of times each track changes direction. (K) Mean absolute acceleration per track. (L-M) Distribution of stationary mitochondria track acceleration summary statistics across both experiments. (L) Mean absolute acceleration per track. (M) Standard deviation of acceleration per track. (N) KymoButler track segmentation for the BR1 experiment, well B04, axon 66, with segmentation errors denoted with arrows.

Our observation above differs from previous work that found decreased mitochondrial motility in *MFN*2^mut^ neurons^15,23,24,45–47^. Those studies differ from ours in model system selected, in the variants profiled, and in annotation strategy (Table 2). To assess whether our choice of automated segmentation tool was responsible for this difference, we manually traced kymographs from wells B07 and F07 of the BR2 experiment, resulting in a total of *n* = 1, 161 manually segmented tracks (Fig. 3d). Manual tracing produced higher motility estimates for both genotypes and resulted in a slightly higher motility percentage in the R364W conditions versus WT (Fig. 3e). Side-by-side inspection showed slight discrepancies, with KymoButler missing smaller motile mitochondria and reporting spurious short-lived stationary mitochondria in kymographs containing large stationary mitochondria (Extended Data Fig. 5a,b).

**Table 2.** Summary of previous publications that performed kymograph-based analysis of mitochondrial motility in *MFN*2^mut^ neurons. Lentivirus (LV) is used to overexpress the mutant *MFN2* allele in model systems of cultured dorsal root ganglion (DRG) neurons.

| Paper | Cell type / model system | *MFN2* variants | Kymograph analysis |
|---|---|---|---|
| Baloh (2007)<sup>45</sup> | LV overexpression, cultured DRG neurons | V69F, L76P, R94Q, P251A, R280H, W740S | manual tracing |
| Misko (2010)<sup>15</sup> | LV overexpression, cultured DRG neurons | R94Q, H361Y | manual tracing |
| Misko (2012)<sup>46</sup> | LV overexpression, cultured DRG neurons | R94Q, R361Y | manual tracing |
| Franco (2020)<sup>47</sup> | reprogrammed CMT2A patient fibroblasts into motor neurons | T105M, R274W, H361Y, R364W | manual tracing |
| Van Lent (2021)<sup>23</sup> | reprogrammed CMT2A patient iPSCs into motor neurons | R94Q | manual tracing |
| Jestice (2026)<sup>24</sup> | reprogrammed edited WT iPSCs into motor neurons | R94Q | KymoButler |
| This study | reprogrammed edited WT iPSCs into motor neurons | R364W, R94W, T105A, W740S | KymoButler & image classifiers |

We next investigated mitochondrial movement metrics beyond the proportion of motile mitochondria, starting with the speed of moving mitochondria, which two studies reported to be slower in *MFN*2^mut^ neurons than in WT neurons^23,47^. In our KymoButler tracks, motile R364W mitochondria were trafficked faster than motile WT mitochondria in the BR2 experiment (Mann-Whitney *p ≤*3.8 *×* 10*^—^*^7^) but slower than WT mitochondria in the BR1 experiment (Mann-Whitney *p ≤*4.8 *×* 10*^—^*^5^; Fig. 3f-g). Weighting motility percentage and track speed by track duration left the direction of both effects unchanged but no longer statistically significant (see Methods; Extended Data Fig. 5c-f). One study reported more pausing among *MFN*2^R94Q^ motile tracks than among WT tracks, with no change in pause duration^24^. We found no difference between WT and R364W in the number of pauses per track, pause duration, or number of direction changes (Fig. 3h-j).

We next computed acceleration-based summary statistics, which, to our knowledge, have not been reported for any mitochondrial kymographs of *MFN*2^mut^ neurons. Mean absolute acceleration was nearly identical between genotypes among motile tracks (Fig. 3k), but stationary R364W tracks had higher mean and higher standard deviation of acceleration than stationary WT tracks (Mann-Whitney *p ≤*4.7 *×* 10*^—^*^27^ and *p ≤*1.3 *×* 10*^—^*^26^; Fig. 3l,m). Stationary R364W mitochondria appear to move back and forth while covering little distance between their start and end time points, which we term “jitter”. On the other hand, stationary WT mitochondria remain still, with no jitter. KymoButler struggles to detect jitter and large mitochondria (Fig. 3n), so we developed alternative approaches for stratifying kymographs between the two genotypes.

### KymoClass robustly distinguishes WT from R364W-mutant kymographs

We next asked whether an image classifier could predict *MFN2* genotype from kymograph images. Since each well contains many axons, and we expect only some axons to display a genotype-associated phenotype, we turned to multiple instance learning (MIL) to train our image classifier^48^. MIL has become a standard approach in computational pathology, where each whole-slide image is divided into thousands of tiles but the diagnosis is considered only for a whole slide in aggregate^49,50^. In attention-based MIL, the model learns a weight for each instance and pools the instances into a single bag embedding using a weighted average^35^. These attention weights indicate which instances the model relied on most for each bag-level prediction^35,50^. We therefore built an attention-based MIL classifier that treats each well as a bag, so that it predicts the genotype of the well and reports which kymographs within the well most influenced that prediction.

We designed a training and evaluation protocol for this MIL classifier. First, we divided each kymograph into overlapping tiles of uniform size (Fig. 4a). Second, we split wells into training, validation, and test sets. We used BR1 wells for training and validation and held out all BR2 wells as a test set, which we evaluated only after training was complete (Fig. 4b). Third, we trained a hierarchical MIL classifier that predicts the genotype of each well from its kymograph tiles (Fig. 4c). The classifier passes each tile through a frozen DINOv3 vision transformer (ViT-L/16) pretrained on the SAT-493M satellite image dataset^34^, which outputs a length-1024 embedding per tile before passing tiles through the learnable encoder, attention pooling blocks, and classification head (see Methods). The model produces two outputs: the predicted genotype of each well and an attention score for each kymograph derived from an axon in that well. We term this kymograph classification approach KymoClass.

**Figure 4.**
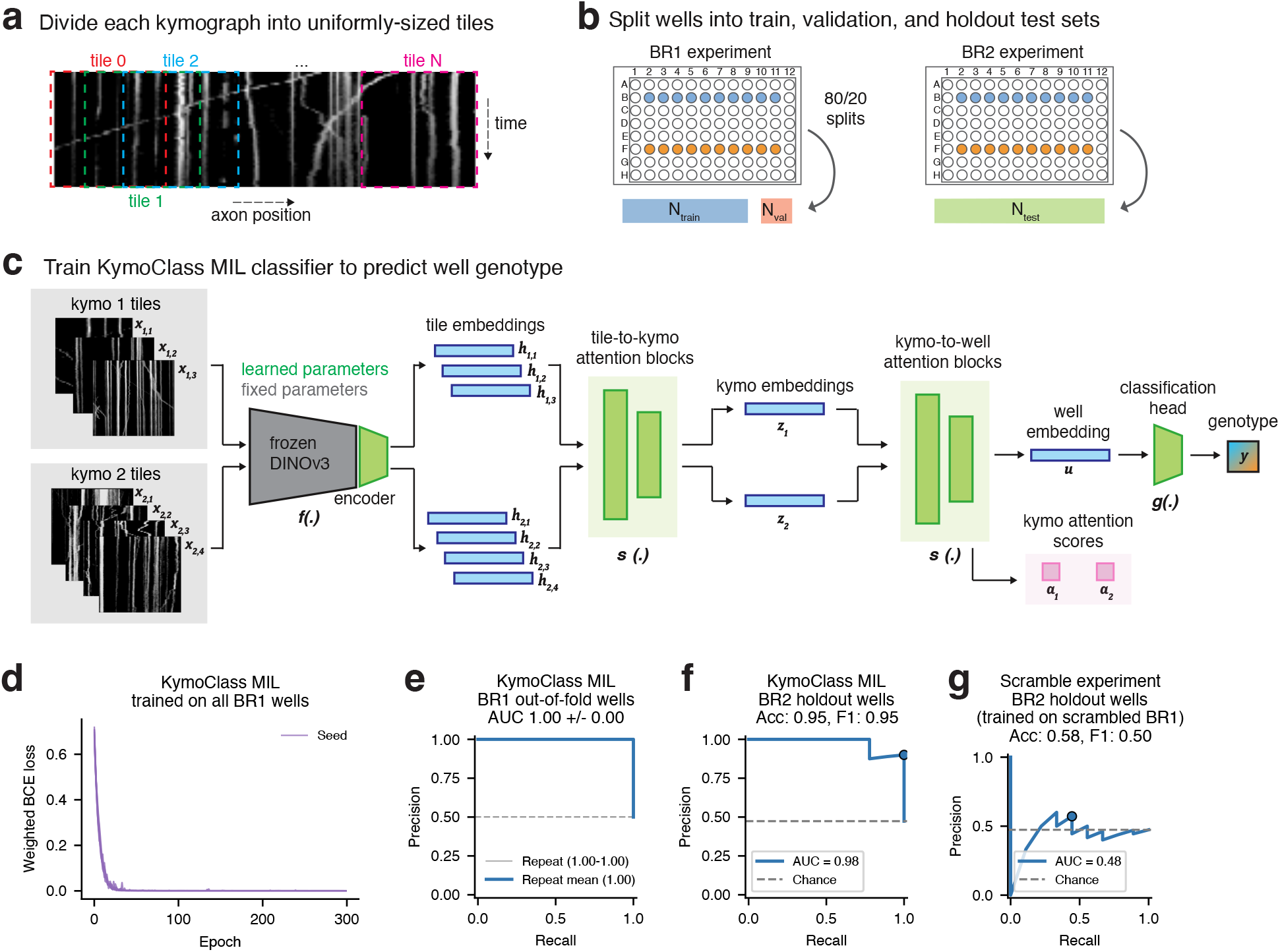
Using KymoClass to predict the genotype of each well. (A-D) Overview of KymoClass model training and evaluation scheme. (A) Each kymograph is divided into uniformly sized, overlapping tiles. Each tile contains all kymograph time points but not all positions of the axon. (B) Wells from the BR1 experiment are used for training and validation, while all wells from the BR2 experiment are held out as a test set. The models in panels D, F, and G are trained on the full BR1 dataset, whereas the cross validation in panel E randomly splits the BR1 wells into train and validation sets using 80/20 splits. (C) KymoClass MIL model diagram. Fixed parameters are shown in grey, learned parameters are shown in green. Intermediate embeddings are shown in blue. (D) Weighted binary cross-entropy (BCE) loss from training KymoClass on all BR1 wells. (E-F) Precision-recall curves for (E) out-of-fold wells in the BR1 train/validation set using 5-fold cross validation and (F) all BR2 wells in the test set. AUC values are noted in the legends. (G) Precision-recall curve for wells in the BR2 test set when KymoClass is trained on BR1 data with scrambled genotype labels. The accuracy and R364W-class F1 scores in all subplot titles represent performance at the 0.5 probability cut, denoted by the single point along each precision-recall curve. The dashed grey line for every precision-recall curve represents random assignment.

We first trained KymoClass on all 20 BR1 wells. The weighted binary cross-entropy loss fell to near zero within the first 50 of 300 training epochs (Fig. 4d). A near-zero training loss shows that KymoClass can distinguish the wells it was trained on, so we assessed its performance on held-out wells in four ways. First, we ran five repeats of 5-fold cross validation over the BR1 wells, with a different fold assignment in each repeat, so that every well was held out once per repeat. In every repeat, KymoClass assigned a higher R364W probability to each held-out R364W well than to any held-out WT well, giving an area under the precision-recall curve (AUC) of 1.00 (Fig. 4e). Second, we applied KymoClass trained on all BR1 wells to the 19 wells of the BR2 test set. It correctly classified all nine R364W wells and nine of the ten WT wells, misclassifying WT well B04 as R364W (accuracy = 0.95, R364W-class F1 = 0.95, AUC = 0.98; Fig. 4f). Third, as a negative control, we randomly permuted the genotype labels across the BR1 wells, trained KymoClass on these scrambled labels, and evaluated it against the true labels of the BR2 test set. This classifier performed at chance level (accuracy = 0.58, R364W-class F1 = 0.50, AUC = 0.48, compared with a chance AUC of 0.47; Fig. 4g). This control shows that the test performance of KymoClass depends on the true genotype labels, and that the model architecture and evaluation procedure alone do not produce high test performance for arbitrary labels. Fourth, as a positive control experiment, we retrained the same classifier with BR2 wells as the training and validation sets and BR1 wells as the test set (Extended Data Fig. 6a). KymoClass trained on BR2 performed comparably to KymoClass trained on BR1, with an out-of-fold AUC of 0.97 over BR2 wells, compared with 1.00 over BR1 wells, and a test AUC of 1.00 on BR1 wells, compared with 0.98 on BR2 wells (Extended Data Fig. 6b,c). This result suggests that our classification strategy forces KymoClass to learn features that discriminate genotype using mitochondrial motion dynamics instead of technical artifacts, such as axon density, that could leak into the kymograph input.

To select the image encoder, and to compare MIL against a simpler baseline, we also trained an instance classifier version of KymoClass. An instance classifier assigns each tile the genotype of its well and predicts genotype one tile at a time (Extended Data Fig. 7a-d). We fit the same logistic regression classifier to the frozen tile embeddings of five pretrained image encoders—two ResNets and three DINO ViTs^34,51–53^—and compared them by 5-fold cross validation over BR1 tiles (see Methods). Mean tile-level AUC ranged from 0.80 *±* 0.02 for ResNet18 to 0.92 *±* 0.01 for DINOv3 SAT-493M, the highest of the five (Extended Data Fig. 7e). We therefore used the DINOv3 SAT-493M encoder in the MIL classifier and in all remaining analyses. With this encoder, the instance classifier predicted tile genotype with accuracy = 0.87 on BR1 validation tiles and accuracy = 0.88 on BR2 test tiles (Extended Data Fig. 7f). Averaging the predicted R364W probabilities of all tiles in a well classified all 19 BR2 wells correctly (Extended Data Fig. 7g). These results demonstrate that well genotype can be accurately predicted by KymoClass without MIL, underscoring that genotype is effectively encoded in nearly every kymograph present in each well.

### MIL attention scores and DINOv3 patch embeddings reveal kymograph properties associated with *MFN2* genotype

After showing that KymoClass was able to accurately and robustly discriminate between healthy and mutant *MFN2* kymographs, we next sought to apply model interpretation techniques to identify image features that contributed to accurate classification. We first visually inspected the kymographs from the BR2 test set that had the highest and lowest attention scores (Fig. 5a,b and Extended Data Fig. 8). High-attention kymographs in WT wells often had straight vertical lines, representative of stationary mitochondria that maintained constant size. High-attention kymographs in R364W wells, by contrast, often had undulating lines consistent with jitter, indicating that mitochondria in these axons rarely maintained a constant velocity or size.

**Figure 5.**
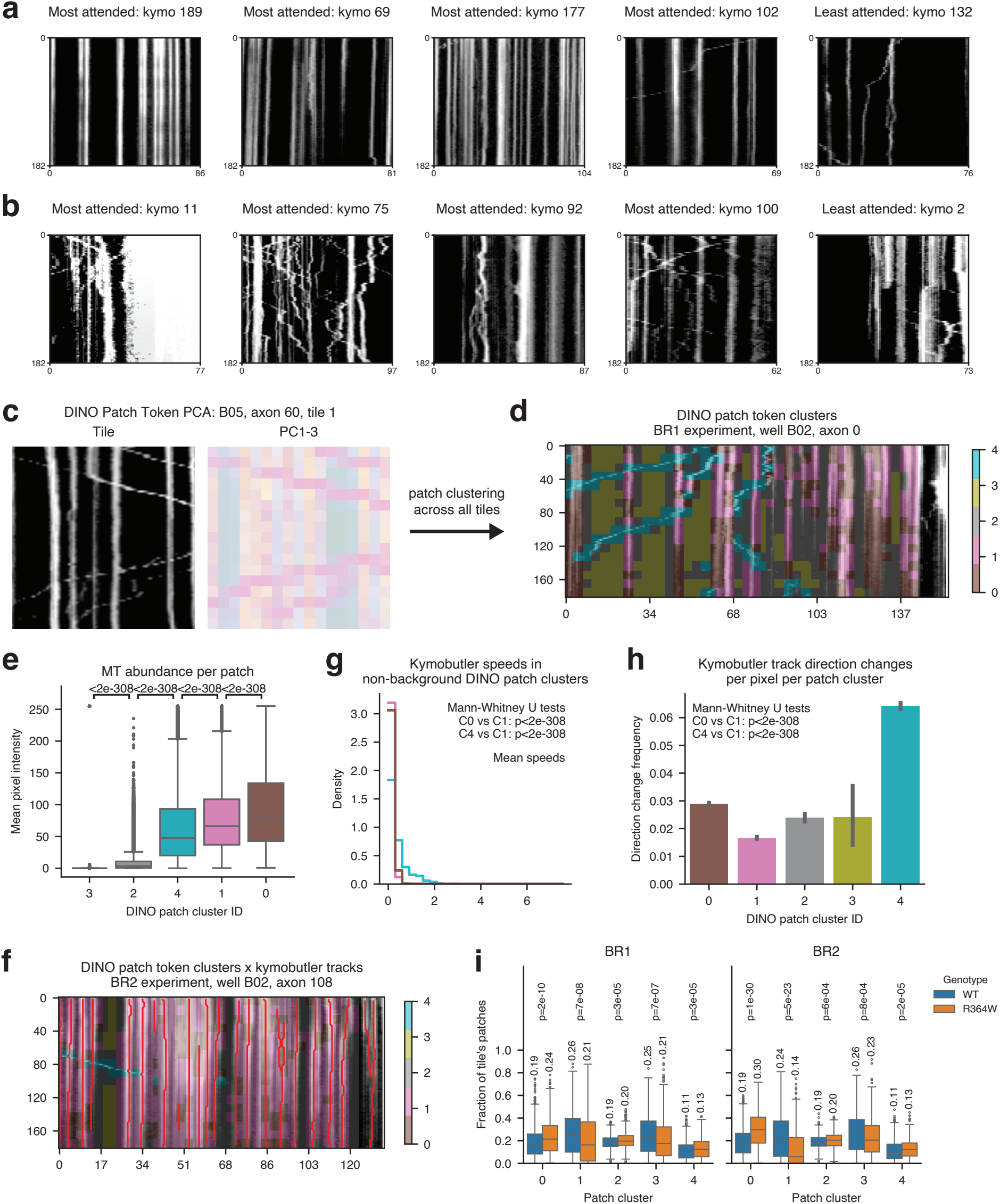
Interpretation of KymoClass model features reveals genotype-to-phenotype relationship based on mitochondrial movement. (A-B) Most and least attended kymographs for (A) WT wells and (B) R364W wells. Each kymograph in the left columns is the most attended kymograph coming from a particular well. The right-most column represents the least attended kymograph from a particular well. (C) DINOv3 patch embeddings top 3 PCs for a representative tile from well B05, axon 60, tile 1. The raw kymograph is shown in black and white on the left. All 3 PCs are shown together in one image using PC1 scores as the red channel, PC2 scores as the green channel, and PC3 scores as the blue channel. (D) Representative example of a kymograph with the cluster ID of each patch indicated by a superimposed color. Patches with no color correspond to areas of the kymograph that had no corresponding tile passed into DINOv3. (E) Mean kymograph pixel intensity, a proxy for mitochondrial abundance, within each 16 *×* 16 DINOv3 patch, split by the DINOv3 patch cluster ID. Clusters are ordered by their mean intensity, and each p-value is from a two-sided Welch’s t-test between the adjacent pair of boxes that it spans, after Benjamini-Hochberg correction across the four comparisons shown. (F) Representative example of a kymograph with the cluster ID of each patch indicated by a superimposed color and all KymoButler tracks superimposed as red lines. (G) Distribution of instantaneous speed for all KymoButler track pixels that appear in a given DINOv3 patch cluster. Only non-background patch clusters are shown. (H) Frequency of KymoButler track direction changes per patch cluster. Each frequency represents the number of direction changes divided by the number of track pixels. Error bars represent 95% confidence intervals from 1000 bootstrap samples. (I) Fraction of each tile’s patches belonging to a given DINOv3 patch cluster, compared between R364W tiles and WT tiles and split by biological replicate. Each box summarizes the tiles of one genotype for one cluster, and that box’s mean fraction is annotated above it. Each p-value is from a two-sided Mann-Whitney U test comparing the tiles of R364W wells against the tiles of WT wells, after Benjamini-Hochberg correction across the ten tests shown in this panel. resolve W740S or T105A from WT.

To systematically interrogate the granular differences between WT and R364W kymographs, we examined the length-1024 patch token embeddings produced for every 16 *×* 16 pixel kymograph patch passed through the frozen DINOv3 encoder^34^. Each patch embedding is a vector that summarizes the visual content of a local patch while also incorporating contextual information from the surrounding patches through the model’s self-attention mechanism, such that visually similar patches can be embedded differently depending on their spatial and temporal context within the larger image. We aggregated all patch embeddings across all wells and experiments into one matrix prior to running principal component analysis (PCA) and K-means clustering (see Methods). PC scores of the first three components visually aligned with stationary mitochondria, moving mitochondria, and background space in kymograph tiles, and the top 50 PCs captured 76% of the total variance in the original length-1024 embeddings (Fig. 5c and Extended Data Fig. 9a-d). The top 50 PCs were used for K-means clustering, and the *K* = 5 results were selected for downstream analysis using the elbow heuristic (see Methods; Extended Data Fig. 9e). Assigning every patch of a kymograph to one of these five clusters produced a semantic segmentation of each kymograph (Fig. 5d).

We next wished to quantitatively define each patch cluster. From visual inspection, we identified two clusters (C0, C1) that correspond to stationary mitochondria, one cluster (C4) that corresponds to moving mitochondria, and two clusters (C2, C3) that correspond to background patches with little-to-no mitochondrial signal (Fig. 5d and Extended Data Fig. 10). To quantitatively determine the amount of mitochondria per patch cluster, we computed the mean pixel intensity within each patch cluster (Fig. 5e). This analysis differentiated the two background clusters C2 and C3; C3 patches have near-zero mitochondrial signal and C2 patches have some mitochondrial signal as they are often the background patches that lie adjacent to mitochondria. Next, we cross-referenced each patch’s cluster ID with the KymoButler tracks that were detected within each patch (Fig. 5f). The number of KymoButler track pixels further confirmed our definition of C2 and C3 being background patches while the others contained mitochondria (Extended Data Fig. 11a). We found that the moving mitochondria cluster C4 had ten-fold faster speed, greater absolute acceleration, a lower fraction of track pixels with 0 *µ*m/s velocity, and more direction changes than the two stationary clusters (Mann-Whitney U test *p ≤*2 *×* 10*^—^*^308^ for all C4 versus C1 tests; Fig. 5g,h and Extended Data Fig. 11b,c). Examining KymoButler metrics similarly enabled us to differentiate between the two clusters of stationary mitochondria, revealing that C0 had approximately two-fold increases in mean speed, mean acceleration, and direction change frequency, along with fewer 0 *µ*m/s velocity track pixels, compared to C1 (Mann-Whitney U test *p ≤*2 *×* 10*^—^*^308^ for all C1 versus C0 tests; Fig. 5g,h and Extended Data Fig. 11b,c). Along with visual inspection, these summary statistics indicated that C0 mitochondria showed jitter, or slight back-and-forth displacement over time, whereas C1 stationary mitochondria were fully stationary.

We next asked whether the abundance of each DINOv3 patch cluster differs between the WT and R364W genotypes. Analyzing the BR1 and BR2 experiments separately, we compared the fraction of each tile’s patches belonging to a given cluster between R364W tiles and WT tiles (Fig. 5i). The two stationary clusters, C0 and C1, had the strongest association with genotype, with C0 “stationary with jitter” patches being enriched in R364W tiles (BR1: *p_ad j_ ≤*2 *×* 10*^—^*^10^; BR2: *p_ad j_ ≤*1 *×* 10*^—^*^30^) and C1 “stationary without jitter” patches being enriched in WT (BR1: *p_ad j_ ≤*7 *×* 10*^—^*^8^; BR2: *p_ad j_ ≤*5 *×* 10*^—^*^23^). Representative time-lapse videos of an R364W axon segment enriched for C0 patches and a WT axon segment enriched for C1 patches are shown in Supplementary Video 2 and Supplementary Video 3, respectively. Background cluster C3 was enriched in WT tiles (BR1: *p_ad j_ ≤*7 *×* 10*^—^*^7^; BR2: *p_ad j_ ≤*8 *×* 10*^—^*^4^) and moving mitochondria cluster C4 was enriched in R364W (BR1: *p_ad j_ ≤*3 *×* 10*^—^*^5^; BR2: *p_ad j_ ≤*2 *×* 10*^—^*^5^); however, these patch clusters were less genotype-specific than C0 and C1. This analysis suggests that the differential abundance of stationary mitochondria subtypes, with and without jitter, is one of the more important features that distinguishes WT from R364W motor neurons.

### KymoClass correctly scores the pathogenicity of all five *MFN2* genotypes

Given that KymoClass robustly discriminated healthy WT from diseased R364W kymographs, we next asked whether the KymoClass model trained on these two genotypes alone could score the mitochondrial trafficking pathogenicity of *MFN2* genotypes that it had never seen during training. To test this, we analyzed time-lapse fluorescence images of iPSC motor neurons carrying three additional *MFN2* mutations—W740S, T105A, and R94W—across the same two biological replicate 96-well plates used above (Fig. 6a; Table 1). We generated kymographs from every axon segment and passed them as input to the previously described WT versus R364W KymoClass MIL model that was trained on BR1, predicting the R364W probability of each well (Fig. 6b; see Methods). Consistent with the reported CMT neuropathy score (CMTNS) severity of these mutations^38^, R94W wells received R364W probabilities as high as those assigned to the R364W wells themselves. On the other hand, W740S and T105A wells received probabilities comparable to those of WT wells. This WT versus R364W KymoClass model therefore separates the two most severe mutations (R364W, R94W) from the two milder ones (T105A, W740S), but it does not

**Figure 6.**
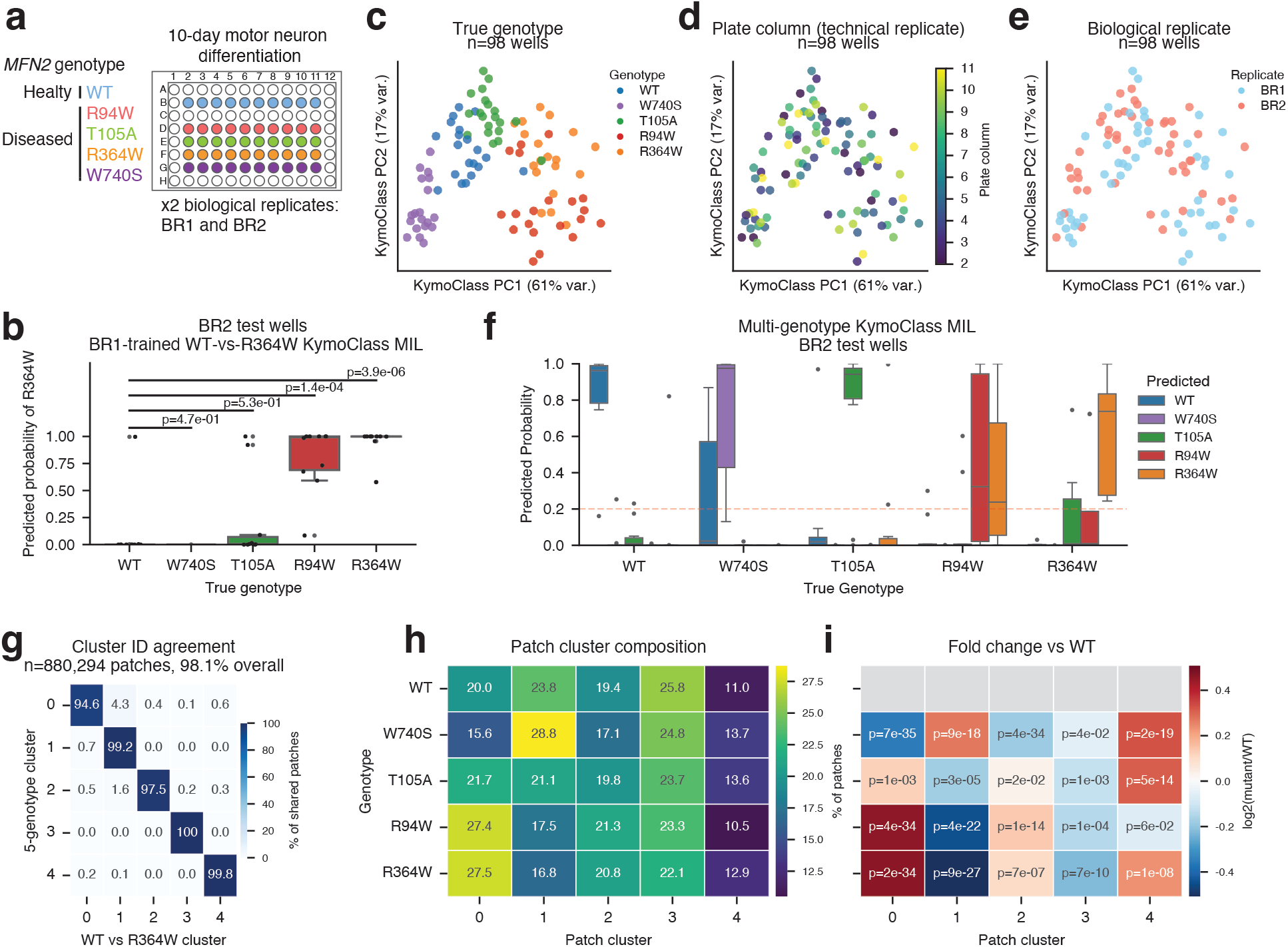
KymoClass correctly characterizes the T105A, W740S, and R94W *MFN2* mutations. (A) Plate map of the WT and *MFN*2^mut^ genotypes, with one 96-well plate per biological replicate. (B) Predicted R364W probability for all wells in the BR2 holdout set, split by true genotype. Probabilities are obtained from the WT versus R364W KymoClass model that was trained using only BR1 wells. P-values are from two-sided t-tests between each mutant genotype versus WT after Benjamini-Hochberg multiple test correction. (C-E) Principal component analysis (PCA) representation of the well embeddings learned by a KymoClass model that was trained on BR1 wells to predict all five genotypes. All *n* = 98 wells of both biological replicates are shown. Each point is a unique well that is colored by its (C) true genotype, (D) plate column, and (E) biological replicate experiment. The percentage of variance explained by each PC is noted in parentheses. (F) Predicted probability that each BR2 holdout well belongs to each of the five genotypes, using a multi-genotype KymoClass model trained on all wells of the BR1 experiment. Boxes are grouped by the true genotype of the well and colored by the predicted genotype. Each box summarizes the technical replicate wells of one true genotype: ten wells each for WT, R94W, and T105A, and nine each for R364W and W740S. The dashed line at a probability of 0.2 denotes the random chance baseline for five genotypes. (G) Correspondence between the two patch clusterings, computed over the 880,294 patches of the WT and R364W wells that both clusterings contain. Rows are the cluster assigned when all five genotypes are clustered together, columns are the cluster assigned when only WT and R364W are clustered, and each cell is the percentage of that row’s patches falling in that column, so rows sum to 100. The diagonal is the per-cluster agreement between the two clusterings. (H) Patch cluster composition of each genotype, given as the percentage of that genotype’s patches assigned to each cluster, so rows sum to 100. (I) The same compositions expressed relative to WT. Color is the log-2 fold change of a mutant genotype’s percentage over the WT percentage, so red marks clusters enriched in the mutant genotype and blue marks clusters depleted in it. The WT row is greyed out because WT is the reference. Each annotated p-value is from a two-sided Mann-Whitney U test of the per-tile fraction of patches in that cluster, mutant tiles versus WT tiles with both biological replicates pooled, after Benjamini-Hochberg correction across the 20 tests shown in this panel. Panels H and I are computed over patches, whereas the p-values in panel I come from a test over tiles.

We next trained a multi-genotype KymoClass model that used the same MIL model architecture as in Fig. 4c, except with a classification head that predicted all five genotype class labels (see Methods). Visualizing KymoClass’s learned embeddings in two dimensions with PCA, we found that wells from the same genotype tended to occupy similar regions of the embedding space (Fig. 6c), whereas plate column and biological replicate labels were dispersed throughout the embedding (Fig. 6d,e), suggesting that KymoClass’s encoder and attention blocks learn genotype-specific embeddings prior to the classification head. When examining predicted genotype probabilities of the BR2 test wells, the multi-genotype classifier showed substantial confusion between the R364W and R94W genotypes, suggesting that these two mutations produce similar mitochondrial trafficking phenotypes (Fig. 6f). This confusion is notable because R364W and R94W carry the two most severe CMT neuropathy scores among the four mutations studied here, yet they alter different domains of the MFN2 protein: R364W lies in the helical domain and R94W lies in the hinge region of the dynamin GTPase domain (Table 1). Although the WT versus R364W KymoClass model scored T105A and W740S wells as WT-like, the multi-genotype KymoClass model cleanly distinguished both genotypes from WT (Fig. 6b,f). These two mutations span mild to moderate clinical severity and lie in different regions of the protein, T105A in the binding region of the dynamin GTPase domain and W740S in the carboxyl terminus (Table 1), so each may confer a trafficking phenotype that the multi-genotype KymoClass model separates from WT along its own axis. As in the WT versus R364W KymoClass model, attention was concentrated on a few kymographs per well: across the five genotypes, the most attended kymograph of a well carried 2.1 to 7.6 times the weight of a uniform average, whereas the least attended kymograph of that well carried 0.4 times the uniform weight or less (Extended Data Fig. 12).

Finally, we computed the DINOv3 patch clusters using kymograph patches from all five genotypes. K-means clustering of this larger set of patches recovered the clusters that we had defined on WT and R364W patches alone: after matching cluster labels between the two clusterings, 98.1% of the 880,294 patches shared by both analyses carried the same label, and the per-cluster agreement ranged from 94.6% to 99.99% (Fig. 6g). We therefore carried the cluster interpretations of Table 3 over to all five genotypes, and compared each mutant genotype’s patch cluster composition against that of WT to ask which mutations share the WT proportions of background, stationary, and moving patches (Fig. 6h,i and Extended Data Fig. 13). Relative to WT, R364W and R94W were enriched to a similar degree in C0 “stationary with jitter” patches (R364W: *p_ad j_ ≤*2 *×* 10*^—^*^34^; R94W: *p_ad j_ ≤*4 *×* 10*^—^*^34^) and depleted in C1 “stationary without jitter” patches (R364W: *p_ad j_ ≤*9 *×* 10*^—^*^27^; R94W: *p_ad j_ ≤*4 *×* 10*^—^*^22^). T105A showed a weaker enrichment of C0 patches (*p_ad j_ ≤*1 *×* 10*^—^*^3^) and a weaker depletion of C1 patches (*p_ad j_ ≤*2 *×* 10*^—^*^5^), in the same direction as R364W and R94W; its largest difference from WT, however, was an enrichment of C4 “moving mitochondria” patches (*p_ad j_ ≤*5 *×* 10*^—^*^14^). W740S ran counter to the other three mutations: relative to WT, it was depleted in C0 patches (*p_ad j_ ≤*7 *×* 10*^—^*^35^) and enriched in C1 patches (*p_ad j_ ≤*9 *×* 10*^—^*^18^). These results suggest that W740S might have a phenotype distinct from the axis that separates WT from R364W.

**Table 3.**
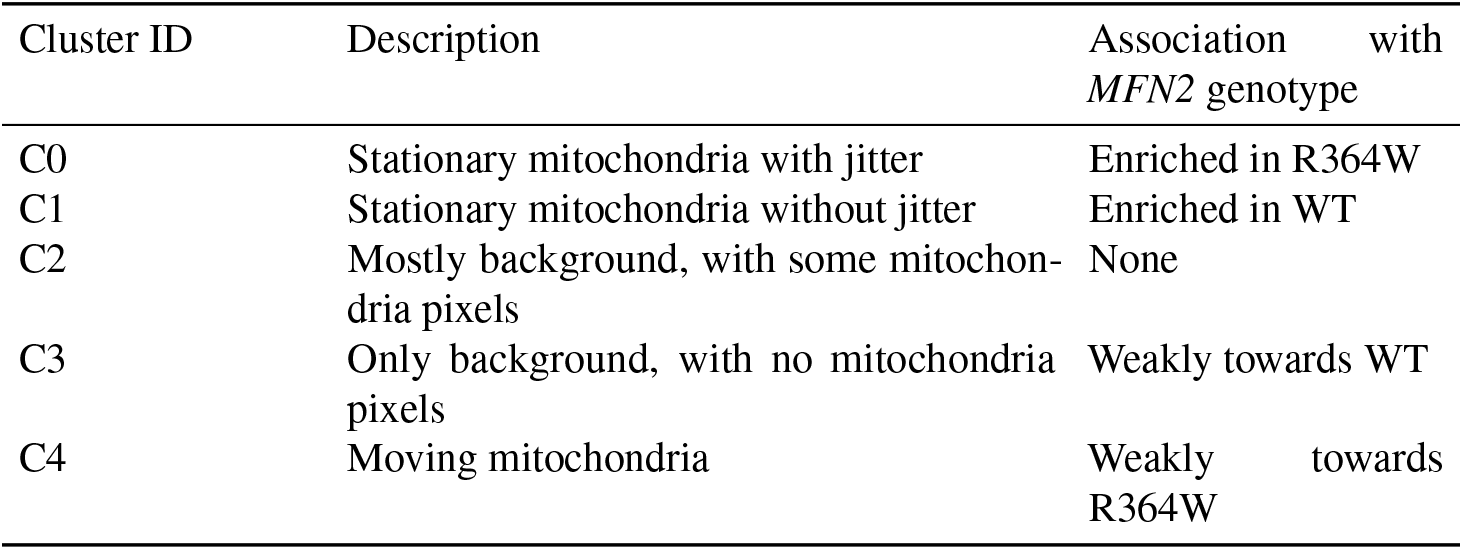
Summary of DINOv3 patch cluster descriptions and their association with *MFN2* genotype. The third column summarizes which genotype each cluster is more abundant in, based on the per-tile cluster fractions compared in Fig. 5i.

| Cluster ID | Description | Association with *MFN2* genotype |
|---|---|---|
| C0 | Stationary mitochondria with jitter | Enriched in R364W |
| C1 | Stationary mitochondria without jitter | Enriched in WT |
| C2 | Mostly background, with some mitochon-dria pixels | None |
| C3 | Only background, with no mitochondria pixels | Weakly towards WT |
| C4 | Moving mitochondria | Weakly towards R364W |

## Discussion

In the 20 years since *MFN2* mutations were first linked to CMT2A, over 100 distinct dominant-negative mutations have been identified. Great effort has been made by clinicians to develop a validated composite scoring system to classify CMT2A disease severity as mild, moderate, or severe. Several cohort studies have applied these CMTESv2 or CMTNS scores across mutation carriers to help build a clearer picture of *MFN2* genotype-phenotype correlations^54,55^. However, no such system exists to score and classify cell pathology across *MFN2* mutations *in vitro*. As technology has advanced, the dissection of *MFN2*-related phenotypes has moved from fibroblasts and mouse models to more physiologically relevant human iPSC-based platforms. Across these systems, the phenotype most consistently associated with mutations in *MFN2* has been dysfunctional mitochondrial motion.

To interrogate this phenotype across *MFN2* mutations, we first engineered an isogenic series of *MFN2* mutant iPSC lines representing severe (R364W, R94W) and milder (T105A, W740S) disease. This strategy allows for greater confidence in *MFN2* mutation-related findings, as genetic background can influence pluripotency and differentiation potential into neurons. Additionally, other variants in patient genomes may influence disease severity, adding unwanted complexity to comparisons of *MFN2*-specific biology. Next, we developed computational approaches for predicting CMT2A-associated mitochondrial motility phenotypes from videos of *MFN*2^WT^ and *MFN*2^mut^ iPSC-derived motor neurons using an MIL platform. Our work highlights the benefits and challenges of using kymographs for analyzing mitochondrial trafficking along axons. We demonstrate that, while manual segmentation of kymographs to obtain individual mitochondrial tracks produces interpretable features, such as speed and pausing, these features do not distinguish *MFN*2^mut^ from *MFN*2^WT^ samples. Additionally, we show that automated kymograph segmentation programs, such as KymoButler, can overestimate or underestimate mitochondrial motion due to small, erratic, or large fused mitochondrial tracks. To address this shortcoming, we developed KymoClass, an image classification approach that leads to highly accurate genotype prediction and unsupervised identification of phenotypes that separate *MFN*2^mut^ from *MFN*2^WT^ samples.

A central biological insight from this work is identifying a subtype of stationary mitochondria with jitter. We found that jitter was most prevalent in *MFN*2^R364W^ and *MFN*2^R94W^ genotypes—the genotypes with the highest reported CMT2A clinical severity. While orthogonal studies are needed to confirm mitochondrial jitter as the phenotype most strongly associated with CMT2A pathogenesis, we hypothesize that jitter could represent a motion pattern that reveals phenotypes related to MFN2’s established role in mediating outer mitochondrial membrane fusion through the formation of MFN1-MFN2 heterodimers and MFN2-MFN2 homodimers^19,56–58^.

Overlaying our data with the biochemical characterization from Li *et al.*^36^, we find that our KymoClass-based groupings map onto the underlying differences in MFN2 GTP hydrolysis and dimerization capacity. Wild-type MFN2 hydrolyzes GTP slowly, forming stable homodimers and heterodimers with MFN1, which allows for tethering of adjacent mitochondrial outer membranes and subsequent fusion. R94W and R364W cluster together in this study, defined by the largest fraction of mitochondrial jitter patches, and represent a clear, severe, dominant-negative disease. Both mutants show rapid GTP turnover while retaining the ability to heterodimerize with WT MFN2 and MFN1, thereby hijacking wild-type proteins into non-productive complexes. R94W differs from R364W in its inability to homodimerize, restricting interactions with the wild-type protein. W740S shows GTP turnover comparable to WT MFN2 and retains its ability to interact with WT MFN2, MFN1, and self-dimerize, consistent with its position outside of the functional GTPase region. In our system, W740S produced less jitter than R94W and R364W, with the highest abundance of fully stationary mitochondria patches (C1), representing a complete arrest of motion. This suggests a dominant-negative mechanism distinct from R94W and R364W. Interpreting our data using the T105A mutant alongside the biochemical characterization of T105M, mutations at this residue are highly consistent with biochemical loss-of-function. In Li’s study, T105M completely loses the ability to bind to WT MFN2, MFN1, or self-dimerize. In this manuscript, the motion patterns of T105A mutants are distinct from those of the three other mutants and wild-type, separating from other conditions when the multi-genotype KymoClass model is applied. The distinct mitochondrial motion patterns observed for each mutation reinforce prior findings linking mutation domain and function to divergent mitochondrial trafficking patterns and downstream clinical phenotype.

Taken together, our results demonstrate the value of KymoClass to identify mitochondrial motion-derived traits. Mutations that may appear indistinguishable from wild-type according to track segmentation summary statistics, such as velocity and motility percentage, instead show clearly distinct phenotypes that only KymoClass can detect. This platform offers a route toward classifying variants at the cellular level. We hypothesize that pathogenic *MFN2* mutations alter mitochondrial motion patterns in both moving and non-moving populations, and that these patterns provide mechanistic insight into a variant’s mechanism of action and severity.

Several limitations warrant consideration. Deep learning image classifiers can learn unknown batch effects, especially when a pretrained model is treated as a black-box feature extractor. We mitigated this risk by evaluating performance across independent differentiation and imaging experiments; however, variation in illumination, staining intensity, or culture conditions could still influence predictions. Expression of both early and mature motor neuron markers confirmed that our cells had fully differentiated into motor neurons by the day of imaging, but neuronal age continues to shape mitochondrial trafficking, and motility deficits in *MFN*2^mut^ neurons may deepen in neurons cultured well past this point. It is possible that neuronal age, not downstream analysis choices, is the reason why our track-derived statistics diverge from the reduced motility of *MFN*2^mut^ mitochondria reported elsewhere^15,23,24,45–47^, but only a matched differentiation time course experiment would test this explanation directly. The phenotypes that KymoClass learned are therefore those present at this stage of differentiation, and users applying KymoClass under different differentiation or preprocessing protocols should retrain or fine-tune the model with their own data.

Looking forward, our results illustrate how establishing quantitative, automated phenotyping criteria enables scalable analysis of organelle movement in iPSC-derived neurons. We view two applications as the most immediate, and a third as a potential future direction. The first application is the characterization of *MFN2* variants of unknown significance^18,59–61^. Based on our results in Fig. 6, we believe that a KymoClass model trained on a small set of well-characterized genotypes can score the pathogenicity of a new variant without that variant appearing anywhere in its training data. The second application is high-throughput therapeutics screening, where the KymoClass pathogenicity score can serve as a per-well readout of phenotypic rescue by a particular therapeutic. Lastly, we believe KymoClass could be relevant for scoring mitochondrial motion in other neurologic conditions. Mitochondrial dysfunction has been reported in Parkinson’s disease^62^, Alzheimer’s disease^63^, and amyotrophic lateral sclerosis^64^. Together, these applications point toward a mode of disease modeling in which we extract and interpret motion-based phenotypes in a high-throughput, data-driven manner—an approach we expect to be valuable well beyond CMT2A.

## Methods

### hiPSC culture

Human iPSCs were cultured on Matrigel (0.337 mg/mL) (Corning)-coated cell culture plates at 37°C and 5% CO_2_, in mTeSR Plus (Stem Cell Technologies) medium, which was changed every other day. Cells were passaged when 60-80% confluent using Accutase and seeded in media containing 10 *µ*M Y-27632.

### Mutation engineering and genotyping

The R364W mutation was engineered into the WTD iPSC line^40^ containing a doxycycline-inducible vector expressing human transcription factors *NGN2*, *ISL1*, and *LHX3* (hNIL) in the *CLYBL* safe harbor locus as previously described^65^.

CRISPR gRNA was designed to target *MFN2* Exon 11 with the sequence 5’-GTTTGAGCAGCACACGGTCC-3’. We used dual 60 bp oligos, one containing the WT sequence with a silent mutation and one containing the mutation sequence, in order to enhance heterozygous insertion events (Extended Data Fig. 2). To prepare the ribonucleoprotein (RNP) complex, 240 pmol of gRNA (IDT) and 120 pmol HiFi SpCas9 were mixed and incubated at room temperature for 20 minutes. 1 *×* 10^6^ iPSCs were transfected with RNP and the homology-directed repair (HDR) template oligos (IDT) using the P3 Primary Cell 4D-Nucleofector X Kit S (Lonza) with pulse code CA137. After nucleofection, cells were immediately collected in mTeSR Plus media containing 10 *µ*M Y-27632 and seeded into 2 wells of a 6-well plate. After 48 hours of culture, single-cell clones were sorted using a BD FACSAria Fusion. Genotypes were verified via Next Generation Sequencing and Sanger Sequencing (for visualization).

### Motor neuron differentiation

WT and R364W-containing iPSCs were differentiated into lower motor neurons as described in^39^, with the following modifications. On day 3, cells were seeded at 20,000 cells per well onto poly-D-lysine (PDL) (Sigma, P7405) and laminin-coated clear bottom imaging 96-well plates. On day 4, the media was removed and replaced with fresh Neural Induction Media (NIM) supplemented with B-27 (Gibco, 17504-044), CultureOne (Gibco, A33202-01), 1 *µ*g/mL laminin, 20 ng/mL BDNF (PeproTech, 450-02), 20 ng/mL GDNF (PeproTech, 450-10), and 10 ng/mL NT3 (PeproTech, 450-03). On day 7, a half volume of media was aspirated and replaced with fresh NIM supplemented with B-27, CultureOne, 1 *µ*g/mL laminin, 40 ng/mL BDNF, 40 ng/mL GDNF, and 20 ng/mL NT3. On day 10, a half volume of media was aspirated and replaced with fresh Neural Maintenance Media (NMM) supplemented with B-27, CultureOne, 1 *µ*g/mL laminin, 40 ng/mL BDNF, 40 ng/mL GDNF, and 20 ng/mL NT3.

### Immunofluorescent staining and imaging

Cells were fixed with 4% PFA, washed twice with PBS, then incubated with 5% Normal Goat Serum (Thermo Fisher Scientific) at RT for 1 hour. After removing blocking buffer and washing twice with PBS, primary antibody was added in 3% NGS and incubated overnight at 4°C. After removing this solution and washing 3 times with PBS, secondary antibodies and DAPI were added in 3% NGS and incubated for 1 hour at RT in the dark. Images were taken on a Keyence BZ-9000 fluorescence microscope with a 40*×* objective or the automated Cell Insight CX7 LZR Pro HCS Platform at 20*×* (Thermo Fisher Scientific). The CX7 microscope has a field size of 444.54 *×* 444.54 *µ*m, and the camera acquisition mode is 1,104 *×* 1,104 (2 *×* 2 binning).

### Gene expression analysis by ddPCR

RNA was extracted from i^3^ lower motor neurons (i^3^LMNs) using the Quick-RNA Miniprep Kit (Zymo Research, catalog no. R1055), according to the manufacturer’s instructions. RNA concentrations were quantified using the NanoDrop spectropho-tometer. Reverse transcription was performed using the iScript cDNA Synthesis Kit (Bio-Rad, catalog no. 1708891). To characterize i^3^LMNs the cDNA (0.5 ng) was amplified in a 25-*µ*L ddPCR reaction using commercially available FAM-labeled TaqMan gene expression assays for *MNX1* (Thermo Fisher, catalog no. Hs00196245_m1) and *CHAT* (Thermo Fisher, catalog no. Hs00193572_m1) in combination with HEX-labeled *GAPDH* gene expression assay (Bio-Rad, catalog no. 10031255) as an internal control for normalization. To measure total *MFN2* mRNA expression a FAM-labeled TaqMan gene expression assay was used in combination with a HEX-labeled *UBE2D2* gene expression assay as an internal control for normalization. To deter-mine allele-specific *MFN2* mRNA expression, i^3^LMN cDNA (0.5 ng) was amplified in a 25-*µ*L ddPCR reaction with TaqMan SNP genotyping assay targeting rs79736124 SNP in the 3’ UTR of *MFN2* (Thermo Fisher, catalog no. C_105316276_10). All ddPCR reactions were analyzed using the Bio-Rad QX200 System and QuantaSoft software. For total mRNA expression of *MNX1*, *CHAT*, and *MFN2*, the ratio of FAM+ to HEX+ droplets was calculated to normalize to *GAPDH* or *UBE2D2*. Fractional abundance of FAM+ versus VIC+ droplets was used to quantify allele specific *MFN2* expression.

### Videos of mitochondrial movement

To visualize axons and mitochondria, iPSCs were transduced with lentivirus from the L304-EGFP-Tubulin-WT (Addgene, 64060) and EF1a-mito-dsRED2 (Addgene, 174541) plasmids, then purified using fluorescence-activated cell sorting (FACS). Reporter iPSCs were differentiated into motor neurons as described above. Motor neurons were imaged on day 10 (BR1 experiment) or day 11 (BR2 experiment) of differentiation using the ImageXpress Micro Confocal Microscope (Molecular Devices). Images were taken every 2 seconds for 3 minutes across 3 z-stacks.

### Video preprocessing

We processed two-channel videos of mitochondrial traffic in cultured motor neurons to remove experimental noise and create kymographs. The red channel measured mitochondria while the green channel measured tubulin expressed in axons and nuclear bodies.

Prior to kymograph construction, we removed technical artifacts from the imaging data represented as xyt volumes. We used contrast-limited adaptive histogram equalization (CLAHE) to correct uneven illumination in the mitochondrial channel^66^. We then clipped the CLAHE-processed mitochondrial volumes using the 5th and 99th percentile values and normalized them between 0 and 255. The clipping removed low-intensity autofluorescence, a common type of noise in 96-well imaging experiments.

### Axon segmentation

After preprocessing the two-channel videos, we then segmented axons from the axon (green) channel. We first maximum-projected the axon volumes along the time axis and applied adaptive histogram equalization (AHE) to remove uneven illumination artifacts from these projections^67^. We then processed the resulting images with a multiscale ridge detection algorithm that incorporated Steger’s, Sato’s, and Lindeberg’s methods^68–70^. Due to variable lentiviral transduction, some axons had particularly faint fluorescence. Therefore, an over-segmentation approach was used to capture all axons. From these processed images, we extracted lines corresponding to the midline of the axons in each well, creating a skeletonized axon mask. False positive axon segmentations tended to be short and thus were filtered out using a ridge contour arc length threshold of 30 pixels. Finally, we joined the skeletonized axons using a binary closing operation. Frequent intersections between axons prevented us from fully segmenting each individual axon.

### Kymograph generation

We used the signal from the mitochondrial channel along the axon segments to create kymographs. We started by converting skeletonized axons into a graph data structure. In this graph, each edge represents an axon fragment, defined by a collection of ordered (*X,Y* ) coordinates. Nodes correspond to axon-axon intersections. Axon fragments consisting of 20 or fewer skeleton points were discarded at this stage. For each remaining axon fragment (edge), we iterated through its ordered (*X,Y* ) coordinates. At each time frame, from first to last, we convolved the mitochondrial volume with a normalized 2 *×* 2 Gaussian kernel (*σ* = 1 pixel) and recorded the smoothed value at each (*X,Y* ) location, so that every kymograph entry is a Gaussian-weighted mean of the mitochondrial signal in the immediate neighborhood of that axon coordinate. We appended these values row-wise to a kymograph array with dimensions corresponding to position within the axon fragment (x-axis) and time frame (y-axis).

We repeated this process for all axon fragments in each well. Finally, we organized the resulting kymograph images by their mutational status and experiment ID. We applied a minimum file size threshold of 1 KB to filter out kymographs from axons that contained insufficient mitochondrial signal. We limited downstream analysis to kymographs that were *>* 62 *µ*m in length.

### Mitochondrial track segmentation with KymoButler

Individual mitochondrial tracks were segmented from kymographs by running KymoButler^31^. We used KymoButler’s bidirectional detection function to segment individual tracks. The only modification we made to the original KymoButler function was disabling color negation during kymograph preprocessing. Disabling color negation appropriately fixed the background as black and mitochondria as white in all kymographs. KymoButler tracks are all 1-pixel-wide lines with integer-valued coordinates. The KymoButler code modification and a Mathematica notebook used for running KymoButler can be found in this forked repository https://github.com/bee-hive/KymoButler.

### Summary statistics of segmented mitochondrial tracks

After segmenting individual mitochondrial tracks within a kymograph, we computed several statistics for each track. Throughout, kymograph coordinates were converted into physical units using an imaging resolution of 0.684132 *µ*m per pixel along the position axis and a frame interval of 2 seconds along the time axis. To compute the speed and duration of each mitochondrial track, we start by defining the xy-coordinates of the first and last point within the track (*X*_0_*,Y*_0_) and (*X_—_*_1_*,Y_—_*_1_), respectively. The track duration (Δ) is the total time duration that a given track appears for

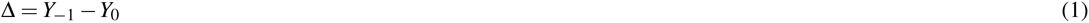

in units of seconds whereas the mean speed (|*θ* |) is defined by the absolute slope between the two points

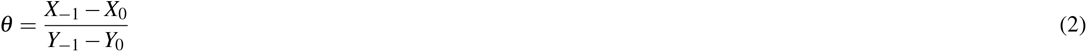

in units of 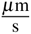.

After computing mean speed and track duration, we computed two secondary summary statistics that used speed and duration as input. The first statistic was the fraction of motile tracks per well (*p*), calculated by thresholding individual tracks into moving or stationary states based on mean speed of 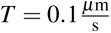

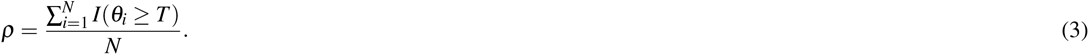

We also created a track-duration adjusted version of the fraction of motile tracks (*p*_!_), giving higher weight to wells in which the moving tracks account for a larger share of the total track duration. We first define the number of moving and stationary tracks in a well, *N*^mov^ and *N*^sta^, along with the cumulative duration of the moving and stationary tracks,

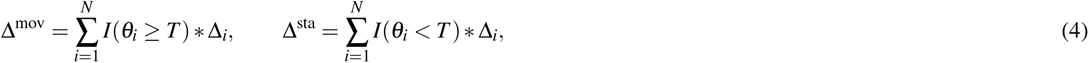

from which we define the duration-weighting factor for moving tracks

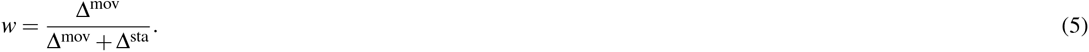

The duration-adjusted fraction of motile tracks then reweights the moving and stationary track counts by *w* and 1 *— w*, respectively,

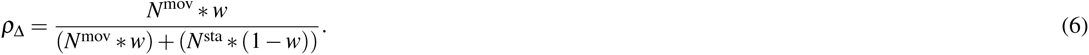

The fraction of motile tracks was used to train a logistic regression genotype classifier. The classifier was trained on the *p* values and genotype labels from the BR1 experiment wells. The classifier was evaluated by running the trained model on BR2 wells and comparing the predicted genotype to the true genotype.

In addition to mean speed and duration of each track that was computed using the start and end coordinates of each track, we computed several pixel-level summary statistics that used the full set of coordinates for each track (*X*_0_*,Y*_0_), (*X*_1_*,Y*_1_)*, …,* (*X_—_*_1_*,Y_—_*_1_). We first were able to compute the instantaneous velocity at each time point *t*

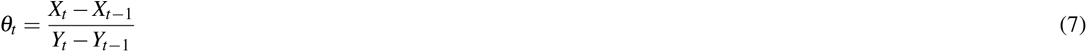

in units of 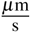 . We were therefore also able to compute instantaneous acceleration in units of 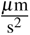 from instantaneous speeds using the following equation:

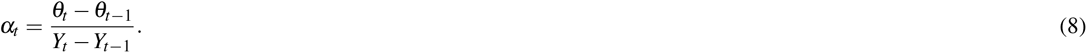

From these per-time point values we computed three per-track summary statistics. First, the mean absolute acceleration per track was computed as the mean of *|α_t_|*, while the standard deviation of acceleration per track was computed as the standard deviation of the signed values *α_t_*. Second, a pause was defined as a consecutive stretch of time points in which *θ_t_* = 0, with the duration of each pause given by the number of time points in that stretch multiplied by the 2 second frame interval; we counted only pauses lasting at least 6 seconds, and the average pause duration per track was computed over these 6 or more second pauses. Note that pauses are defined by exactly zero displacement between consecutive time points rather than by the 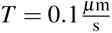 threshold used to stratify whole tracks into moving and stationary states. Third, the number of direction changes per track was computed as the number of times that *θ_t_* changes sign, after first removing all time points at which *θ_t_*= 0, so that a pause between two movements in the same direction is not counted as a direction change and a reversal interrupted by a pause is counted only once.

### Kymograph preprocessing for input into KymoClass

We employed additional filtering and preprocessing steps before passing kymograph images into an image classifier. We only used kymographs with a length greater than 62 *µ*m (91 pixels) and a file size greater than 2 KB (using a PNG encoding). This thresholding removes small axon fragments, likely noise artifacts, that get created during axon segmentation. Likewise, axon fragments that do not contain any mitochondria (all black) fall below 2 KB in file size and thus we remove these kymographs as it would be impossible to determine the genotype from empty kymographs. The remaining kymographs were then upsampled by a factor of three using bicubic interpolation. After upsampling, kymographs were split into uniformly sized overlapping tiles of 272 *×* 272 pixels with a stride of 90 pixels, tiling from left to right. Within each valid kymograph, we only analyzed tiles with a pixel intensity standard deviation of at least 25. Removing low-variance tiles was essential as they were also likely to be empty.

### KymoClass to predict WT versus R364W genotype

We trained two families of KymoClass image classifiers on these kymograph tiles, differing in which object carries the genotype label. A KymoClass multiple instance learning (MIL) classifier treats each well as a labeled bag of unlabeled tiles and predicts one genotype per well. A KymoClass instance classifier instead assigns every tile the genotype of the well it came from and predicts one genotype per tile. KymoClass uses the MIL architecture by default. Both classifiers read the same input: the length-1024 `cls` token that the frozen DINOv3 ViT-L/16 encoder pretrained on SAT-493M returns for each 272 *×* 272 pixel tile^34^, selected by the encoder comparison described in the next subsection. These two classifiers, and all of the patch embedding analyses that follow, used only tiles from the WT and R364W wells of each plate; the three additional *MFN*2^mut^ genotypes present on the same plates were held aside and analyzed separately. All model fitting used scikit-learn v1.7.2^71^ or PyTorch v2.5.1 with torchvision v0.20.1^72^.

#### Multiple instance learning classifier

The KymoClass MIL classifier is a hierarchical gated attention network^35^ (Fig. 4c). Consider one well holding *m* kymographs, where kymograph *i* contributes *n_i_* tiles, and write **x***_i, j_* for the *j*th tile of kymograph *i*. An encoder *f* maps every tile to **h***_i, j_* = *f* (**x***_i, j_*) *∈* R^64^. It is the composition of the frozen DINOv3 backbone, which returns the length-1024 `cls` token of a tile, a per-dimension standardization using the means and standard deviations of the training wells’ tiles, and a learned block of dropout with *p* = 0.25, a linear layer R^1024^ *≃* R^64^, a GELU nonlinearity, and a second dropout with *p* = 0.25. Only the learned block is updated during training. Two gated attention blocks then pool these tile embeddings. A gated attention block *s* scores one embedding **h** as

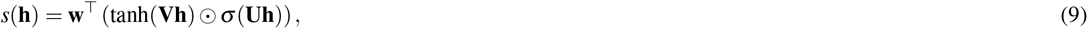

where **V**, **U** *∈* R^32^*^×^*^64^, **w** *∈* R^32^, ʘ is the elementwise product, and *σ* is the logistic function. The first block, *s_α_* , pools the tiles of each kymograph into a kymograph embedding **z***_i_ ∈* R^64^,

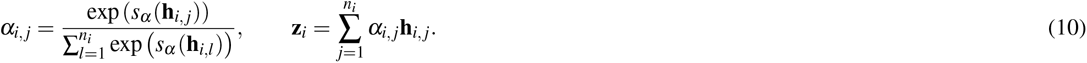

The second block, *s_β_*, holds its own **V**, **U**, and **w** of the same shapes and pools the *m* kymograph embeddings into one well embedding **u** *∈*R^64^,

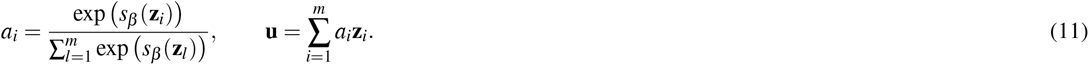

The tile attention weights *α_i, j_* sum to one within a kymograph and the kymograph attention weights *a_i_* sum to one within a well, so both pooled embeddings are convex combinations of their inputs and wells contribute embeddings on the same scale whether they hold few tiles or many. A linear classification head *g* maps the well embedding to the predicted probability that the well carries the R364W mutation, 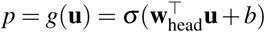. The learned encoder block, the two attention blocks, and the head hold 74,051 learned parameters, 65,600 of which belong to the 1024 *≃* 64 linear layer.

We fit the MIL classifier with AdamW at a learning rate of 10*^—^*^3^, a weight decay of 0.01, and 300 epochs, taking one full-batch gradient step over all training wells per epoch. The loss is binary cross-entropy on the well probabilities, weighting the R364W class by the ratio of WT wells to R364W wells in the training set so that the two genotypes contribute equally at the level at which the labels live. We applied two label-preserving augmentations, redrawn at every epoch. First, each tile entered that epoch’s bag with probability 0.8, so no two epochs saw the same well. Second, each surviving tile was replaced with probability 0.5 by the embedding of its horizontally mirrored crop. Mirroring left to right preserves the label because the x-axis of a kymograph is position along an axon and the traced skeleton starts at an arbitrary end of that axon; we never flipped vertically and never rotated, as either operation would reverse or mix the time axis. Mirrored embeddings were precomputed by passing the mirrored tile crops through the same frozen encoder with the same normalization. A training fold holds at most 20 wells, few enough that one fit’s performance moves with the random seed, so every fit is an ensemble of five models differing only in initialization and augmentation draw (seeds 42 through 46), whose predicted probabilities and attention weights we averaged. Every evaluation used the original tile orientations, with all tiles present and dropout disabled.

We split wells into training, validation, and test sets. The BR1 plate contributed 20 wells (10 WT, 10 R364W) for training and validation, and all 19 wells of the BR2 plate (10 WT, 9 R364W) were held out as a test set that we scored only after training was complete. We evaluated the classifier three ways. First, we drew a single stratified 80/20 split of the BR1 wells (`StratifiedShuffleSplit` with `n_splits=1`, `train_size=0.8`, and `random_state=42`), leaving 16 training and 4 validation wells. Second, and as the primary internal evaluation, we ran five repeats of stratified 5-fold cross validation over the BR1 wells (`StratifiedKFold` with `n_splits=5`, `shuffle=True`, and `random_state` set to 42 plus the repeat index), so every well is held out exactly once per repeat and each repeat draws a different fold assignment. Third, we refit the classifier on all 20 BR1 wells and applied it to all 19 BR2 wells. In each case the per-dimension standardization was fit on the training wells alone. We report the area under the precision-recall curve, computed as average precision, alongside the accuracy and the R364W-class F1 score at a probability threshold of 0.5. We report precision-recall rather than ROC because the number of wells per genotype is unbalanced in several of these evaluations.

#### Instance classifier

The KymoClass instance classifier is a logistic regression fit to the same frozen `cls` token embeddings, without standardization, that predicts the genotype of each tile (0 is WT, 1 is R364W). Fitting a logistic regression to frozen features is equivalent to training a linear probe with a sigmoid activation on top of the encoder. We used scikit-learn’s `LogisticRegression` with an L2 penalty (*C* = 1.0), balanced class weights, the lbfgs solver, and a limit of 1000 iterations. Balanced class weights matter here but not in the MIL classifier, because WT wells contribute roughly 1.7 times as many tiles as R364W wells while each well contributes exactly one bag. The model returns the probability that a tile belongs to the R364W genotype, and we scored a well by averaging that probability over all of its tiles.

We split kymographs, rather than tiles, for the instance classifier. All BR1 kymographs were divided 80/20 into a training and a validation set with `GroupShuffleSplit` (`n_splits=1`, `train_size=0.8`, and `random_state=42`) grouped by (well, kymograph), so that every tile of a kymograph lands on the same side of the split and no kymograph contributes tiles to both sets. Splitting was otherwise unstratified, so both sets contain tiles from every BR1 well. All tiles of the BR2 plate were held out as a test set.

### Testing different image encoder backbones for KymoClass

To determine which pretrained image encoder we should use for our KymoClass MIL classifier, we tested five different KymoClass instance classifiers using the following frozen feature extractors for kymograph tiles: ResNet18 and ResNet50 with supervised ImageNet-1k weights (torchvision IMAGENET1K_V1)^51,73^, DINOv2 ViT-L/14 pretrained on LVD-142M^53^, DINOv3 ViT-L/16 pretrained on LVD-1689M, and the distilled DINOv3 ViT-L/16 pretrained on SAT-493M^34^.

Every encoder expects three-channel input, so each grayscale tile was replicated across three channels. Each tile was then resized to the largest multiple of the encoder’s patch size that does not exceed 272 pixels, which is 266 pixels for the patch-14 DINOv2 model and 272 pixels for the patch-16 ViTs and both ResNets. Tiles were normalized using the channel means and standard deviations of the dataset that the encoder was pretrained on: (0.430, 0.411, 0.296) and (0.213, 0.156, 0.143) for SAT-493M, and (0.485, 0.456, 0.406) and (0.229, 0.224, 0.225) for ImageNet, which we also used for the DINOv2 and LVD-1689M encoders.

For ResNet18 and ResNet50, we replaced the final fully connected layer with an identity map and used the globally average-pooled penultimate features, of length 512 and 2048, respectively. For each ViT, we obtained 1) a length-1024 `cls` token embedding and 2) a 17 *×* 17 *×* 1024 `patch` token embedding tensor per 272 *×* 272 pixel tile. All encoders were run in evaluation mode with gradients disabled, so their features are deterministic given the weights.

We chose among the five encoders by fitting the same logistic regression probe, with the same hyperparameters described above, to each encoder’s frozen tile features and comparing the resulting classifiers. This comparison used tiles from the BR1 experiment only. Kymographs were computed and tiled once, so that every encoder received an identical set of 272 *×* 272 pixel crops.

Evaluation used five-fold cross validation of the BR1 tiles (`StratifiedGroupKFold` with `n_splits=5`, `shuffle=True`, and `random_state=42`), grouped by (well, kymograph) so that no kymograph contributed tiles to both the training folds and the held-out fold, and stratified by genotype. Five folds give each fit the same 80/20 proportion as the instance classifier above, and across the five folds every tile is held out exactly once. The folds were computed once and reused for every encoder, so the comparison is paired. For each fold, the probe was fit on the four training folds and scored on the held-out fold at the tile level, and each fold’s precision-recall curve was interpolated onto a shared grid of 101 evenly spaced recall values. We report the across-fold mean of these interpolated curves, the full range across the five folds, and the mean and standard deviation of the five per-fold AUC values, each computed as average precision from the raw scores rather than from the averaged curve. We report precision-recall rather than ROC because WT tiles outnumber R364W tiles, so a chance classifier reaches a precision equal to the fraction of tiles that are R364W rather than a fixed diagonal. The DINOv3 SAT-493M encoder was selected on the basis of this comparison and used for every subsequent analysis.

### Label scrambling and plate-swap control experiments

We ran two control experiments to assess KymoClass’s performance. Both reused the architecture, the hyperparameters, the splits, and the five-model seed ensemble described above, so each differs from the reported classifier in exactly one respect.

In the negative control experiment, we permuted the genotype labels of the training wells. We drew one uniform random permutation of the vector of BR1 well labels with the numpy default random generator seeded with 0. Permuting a vector of labels rather than drawing new ones preserves the class balance exactly, so the control model trains on the same 10 WT and 10 R364W bags as the reported model. The held-out cross validation folds and every BR2 test well retained their true genotype labels, so each classification task asks whether a model fit to scrambled labels retains any ability to discriminate genotype.

In the positive control experiment, we swapped the roles of the two plates. We trained and cross validated on the 19 BR2 wells and held out the 20 BR1 wells as the test set, leaving every other detail of the procedure unchanged. We then evaluated prediction accuracy for the out-of-fold wells of the training plate and the external test plate. A classifier that had learned a feature specific to one plate, such as axon seeding density, rather than a genotype-associated phenotype should not survive this swap.

### KymoClass attention scores

We used KymoClass’s kymograph attention weights *a_i_*to identify which axons a well’s genotype call rested on, averaging them over the five members of the seed ensemble. Because these weights are normalized within a well, *a_i_* is not comparable between wells that hold different numbers of kymographs: a well of *m* kymographs would give each one a weight of 1*/m* if attention were uniform. We therefore report each kymograph’s attention both as its raw weight *a_i_* and as the ratio *ma_i_* of that weight to the uniform weight of the same well. A high attention weight means that a kymograph shaped the well embedding, not that the kymograph resembles a particular genotype; the genotype call is made by the classification head from the pooled well embedding.

### Visualization and clustering of DINOv3 patch embeddings

We inspected the patch embeddings of WT and R364W kymograph tiles to build intuition for which kymograph properties are captured by the DINOv3 model. As DINOv3 is a ViT model with 16 pixel patch size, a 272 *×* 272 pixel kymograph tile gets broken into a 17 *×* 17 grid of patches where each patch is 16 *×* 16 pixels in size. When passing an image through the DINOv3 model, each 16 *×* 16 patch gets assigned a length-1024 embedding vector. We collated the embedding vectors into a matrix across all 289 patches per tile across all *N*_tiles_ tiles, resulting in a matrix with the shape (*N*_tiles_ *\** 17^2^) *×* 1024. We performed principal component analysis (PCA) on this (*N*_tiles_ *\** 17^2^) *×* 1024 patch embedding matrix to reduce each patch into the top 50 PCs. We clustered the PCA patch embeddings by applying K-means clustering on the (*N*_tiles_ *\** 17^2^) *×* 50 PCA matrix. We performed K-means clustering for values of *K* ranging from 3 to 10. For each *K*, we calculated the inertia value and used these values to create an elbow plot, leading us to select *K* = 5 as the optimal clustering for downstream analysis.

### Differential abundance of DINOv3 patch clusters between WT and R364W tiles

We tested whether each DINOv3 patch cluster is differentially abundant in R364W kymographs relative to WT kymographs. The unit of observation is the kymograph tile rather than the individual patch. Every tile contributes the same 17^2^ = 289 patches, so treating those patches as independent observations would inflate the sample size by a factor of 289 without adding independent information. For each tile *j* and each patch cluster *C*, we computed the fraction of that tile’s patches assigned to *C*,

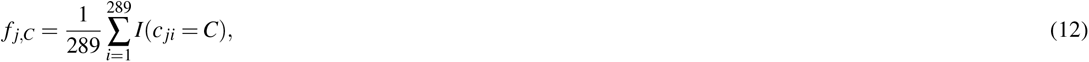

where *c _ji_*is the cluster assigned to the *i*th patch of tile *j* and *I*(*·*) is the indicator function. Each tile therefore contributes one fraction per cluster, and the five fractions of a tile sum to one.

Within each biological replicate, and separately for each patch cluster *C*, we compared the fractions of the R364W tiles against the fractions of the WT tiles,

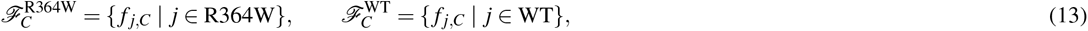

using a two-sided Mann–Whitney U test (`scipy.stats.mannwhitneyu`). We quantified the size of each difference with Cohen’s *d*,

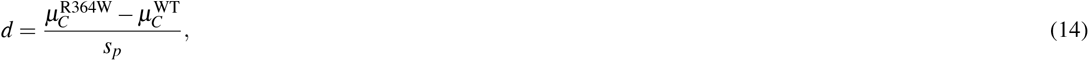

where *µ_C_*^R364W^ and *µ_C_*^WT^ denote the means of *F_C_* ^R364W^ and *F_C_* ^WT^, respectively, and *s_p_* is their pooled standard deviation, so that a positive *d* marks a cluster enriched in R364W tiles.

These tests used the 3,046 WT and R364W tiles that passed the filtering described above: 1,008 WT and 608 R364W tiles from BR1, and 1,006 WT and 424 R364W tiles from BR2, comprising 880,294 patches in total. Testing five clusters within each of the two biological replicates gives ten hypothesis tests, whose p-values were adjusted together for multiple hypothesis testing using the Benjamini-Hochberg false discovery rate (FDR) correction (`statsmodels.stats.multitest.multipletests` with `method=’fdr_bh’`), and adjusted p-values below 0.05 were called significant.

### Quantitative interpretation of DINOv3 patch clusters

To interpret the image-level and dynamic properties captured by each DINOv3 patch cluster, we computed summary statistics based on raw pixel intensities and KymoButler-derived mitochondrial motion features.

As a first approach, we quantified the mean raw pixel intensity of patches within each cluster. For a given patch cluster *C*, let *P_C_* denote the set of all 16 *×* 16 pixel patches assigned to cluster *C*. The mean pixel intensity for cluster *C* was computed as

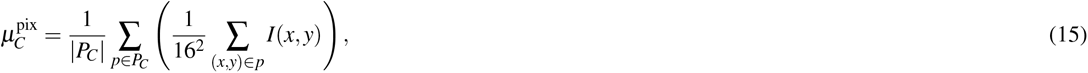

where *I*(*x, y*) denotes the raw pixel intensity at spatial location (*x, y*) in the kymograph image.

As a second approach, we examined the mitochondrial dynamics associated with each patch cluster by intersecting KymoButler track pixels with DINOv3 patches. For each patch cluster *C*, we identified the set of all KymoButler track pixels whose coordinates intersected with patches assigned to *C*, yielding the set

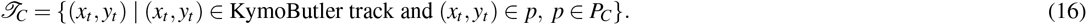

For all pixels in *T_C_*, we aggregated the instantaneous velocities *θ_t_* and absolute accelerations *|α_t_|*, as defined previously. These values were used to compute cluster-level summary statistics, including the fraction of track pixels with zero instantaneous velocity,

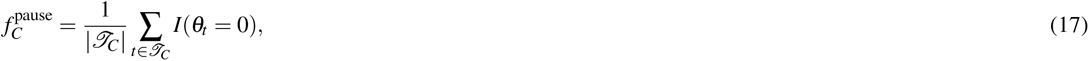

as well as the number of direction changes occurring within cluster *C*, defined as the number of sign changes in *θ_t_* across consecutive time points for track pixels intersecting patches in *P_C_*.

The p-values of these tests are extremely small, even after false discovery rate correction, because the unit of observation is the individual patch. Due to numerical underflow limitations in floating-point arithmetic, p-values reported by Python as 0 *×* 10^0^ were instead reported as *<* 2 *×* 10*^—^*^308^, corresponding to the smallest positive representable floating-point value.

### Scoring additional *MFN2* genotypes with the WT versus R364W KymoClass model

In addition to the WT and R364W rows analyzed above, each plate contained rows of heterozygous R94W, T105A, and W740S motor neurons. The assignment of genotypes to plate rows differed between the BR1 and BR2 experiments. Kymographs from all genotypes were generated, filtered, tiled, and passed through the frozen DINOv3 SAT-493M backbone using the same procedure and parameters described above, yielding one length-1024 `cls` token embedding per tile.

To score the mitochondrial trafficking pathogenicity of the three additional genotypes, we loaded the WT versus R364W KymoClass model that had been trained on all 20 BR1 wells—the same seed ensemble, with the same standardization fit on its own training tiles, that was evaluated on the BR2 test wells above—and applied it unchanged to every well of all five genotypes in both experiments. Each well is one bag, so this gives one predicted R364W probability per well. The WT and R364W wells of BR1 were this model’s training data and are in-sample for it, so every per-genotype comparison was restricted to the 48 wells of the BR2 plate, which the model never saw.

### Multi-genotype KymoClass model

We additionally trained a multi-genotype KymoClass model that predicts which of the five genotypes—WT, R94W, T105A, R364W, or W740S—each well carries. The 50 wells of the BR1 plate were used for training and validation and the 48 wells of the BR2 plate were held out as a test set. The network, its hyperparameters, and its evaluations are those of the WT versus R364W classifier, with two changes. The classification head *g* returns five logits rather than one, and the loss is cross-entropy over the five classes, weighting class *c* by *N/*(5*n_c_*), where *n_c_* is the number of training wells of genotype *c* and *N* is the total number of training wells, so that each genotype’s wells contribute equally. A well’s predicted genotype is the class that receives the highest predicted probability. With five roughly balanced classes, 0.2 is both the uninformative probability and the chance accuracy.

To visualize what the multi-genotype classifier learns below its classification head, we embedded the KymoClass length-64 well embeddings **u** of every well in two dimensions with PCA. These embeddings come from the first member of the seed ensemble of the KymoClass model trained on all BR1 wells, not from the ensemble average: the members differ in initialization and each learns its own arbitrary basis for the length-64 space, so averaging their embeddings would average vectors expressed in different coordinate frames. Both plates were embedded in a single PCA fit so that they share one coordinate system.

### Multi-genotype clustering of DINOv3 patch embeddings

To ask whether the patch clusters defined on WT and R364W kymographs also describe the three additional *MFN2* mutations, we repeated the patch embedding clustering using tiles from all five analyzed genotypes. Kymographs from the WT, R94W, T105A, R364W, and W740S wells of both biological replicates were generated, filtered, tiled, and passed through the frozen DINOv3 SAT-493M backbone using the procedure and parameters described above; the homozygous R94W row was excluded, as in every other analysis reported here. This yielded 7,305 tiles—1,008 WT, 1,086 W740S, 707 T105A, 527 R94W, and 608 R364W tiles from BR1, and 1,006 WT, 804 W740S, 673 T105A, 462 R94W, and 424 R364W tiles from BR2—and 7305 *×* 17^2^ = 2,111,145 patch token embeddings of length 1024.

We pooled the patch embeddings of both biological replicates into a single 2,111,145 *×* 1024 matrix, reduced it to the top 50 principal components, and clustered these components with K-means, using the same procedure and parameters as the two-genotype clustering above. We swept *K* from 3 to 10 and again selected *K* = 5 from the elbow plot. PCA and K-means were fit with scikit-learn v1.7.2^71^ using `random_state=42` and `n_init=10`, without access to genotype or biological replicate labels.

K-means assigns arbitrary integer labels to its clusters, so the labels of this five-genotype clustering are not comparable to the labels of the two-genotype clustering by index. The two clusterings share every patch of the WT and R364W wells, which is 880,294 of the 2,111,145 patches clustered here, so we read the correspondence between them off those shared patches. We tabulated the five-genotype label against the two-genotype label over the shared patches, giving a 5 *×* 5 contingency table, and chose the one-to-one relabeling of the five-genotype clusters that maximizes the total number of patches receiving the same label under both clusterings. This assignment problem was solved with the Hungarian algorithm^74^ (`scipy.optimize.linear_sum_assignment`) applied to the negated contingency table. Under this relabeling, 863,470 of the 880,294 shared patches (98.1%) carry the same cluster label in both clusterings, and the per-cluster agreement ranges from 94.6% for C0 to 99.99% for C3. All five-genotype results are reported with these aligned labels, so C0 through C4 carry the interpretations given in Table 3.

We summarized the patch cluster composition of each genotype as the percentage of that genotype’s patches assigned to each cluster, so that the five percentages of a genotype sum to 100. We report each mutant genotype’s composition relative to WT as the log-2 fold change log_2_(*u_g,C_/u*_WT_*_,C_*), where *u_g,C_* is the percentage of genotype *g*’s patches assigned to cluster *C*.

We tested each mutant genotype against WT for differential cluster abundance using the per-tile fraction *f _j,C_*, the two-sided Mann–Whitney U test, and Cohen’s *d* defined above, substituting the mutant genotype for R364W. We ran this comparison with the two biological replicates pooled and within each replicate separately. These two runs analyze overlapping data to answer different questions, so their p-values were adjusted in two separate Benjamini-Hochberg families: the 20 pooled tests (four mutant genotypes across five clusters) form one family, and the 40 per-replicate tests (four mutant genotypes across five clusters within each of two replicates) form a second. The adjusted p-values reported in the text, in Fig. 6i, and in Extended Data Fig. 13 are those of the pooled family.

The composition percentages and the adjusted p-values summarize the same patches at two different levels: a percentage is computed over patches, whereas a p-value is computed over tiles. A cluster with a large fold change can therefore carry a modest p-value when the per-tile fractions of the two genotypes overlap heavily, and a cluster with a small fold change can carry a small p-value when they do not.

## Data availability

All raw images and processed kymographs are uploaded to BioImage Archive at https://www.ebi.ac.uk/biostudies/bioimages/studies/S-BIAD4158. Processed data (classifier weights, patch clustering results, and statistical test results) will be uploaded to Zenodo under a unique DOI upon publication of this manuscript. DINOv3 model weights can be freely downloaded from Meta at the following site: https://ai.meta.com/research/dinov3/. Contact B.R.C. at for questions regarding iPSC cell lines.

## Code availability

All code used for kymograph generation, KymoClass model training, and downstream analysis can be found in the following GitHub repository https://github.com/bee-hive/conklin_mitochondria_traffick. This repository is currently private but will be made public after final publication. Instructions for installing all KymoClass Python dependencies can be found in the repository’s README.md file. Mathematica code for running KymoButler on our data can be found in our fork of the original KymoButler GitHub repository https://github.com/bee-hive/KymoButler.

## Acknowledgments

We would like to thank all members of B.E.E.’s and B.R.C.’s labs for helpful feedback on this work.

## Author contributions statement

A.C.W., B.L.M., L.E., B.R.C., and B.E.E. conceived the study. B.L.M. and K.R.K. planned and conducted the wet lab experiments. A.C.W. and L.E. planned and conducted the computational experiments. A.C.W., B.L.M., L.E., B.R.C., and B.E.E. analyzed the results and wrote the manuscript. All authors reviewed the manuscript.

## Ethics declarations

### AI usage disclosure

During the preparation of this work, the authors used Claude Code, Google Gemini, and GitHub Copilot to assist with coding tasks. The authors also used Claude and ChatGPT to assist in editing the text of the manuscript. The authors reviewed and edited all content as needed and take full responsibility for the content of the publication.

## Extended Data

**Extended Data Fig. 1.**
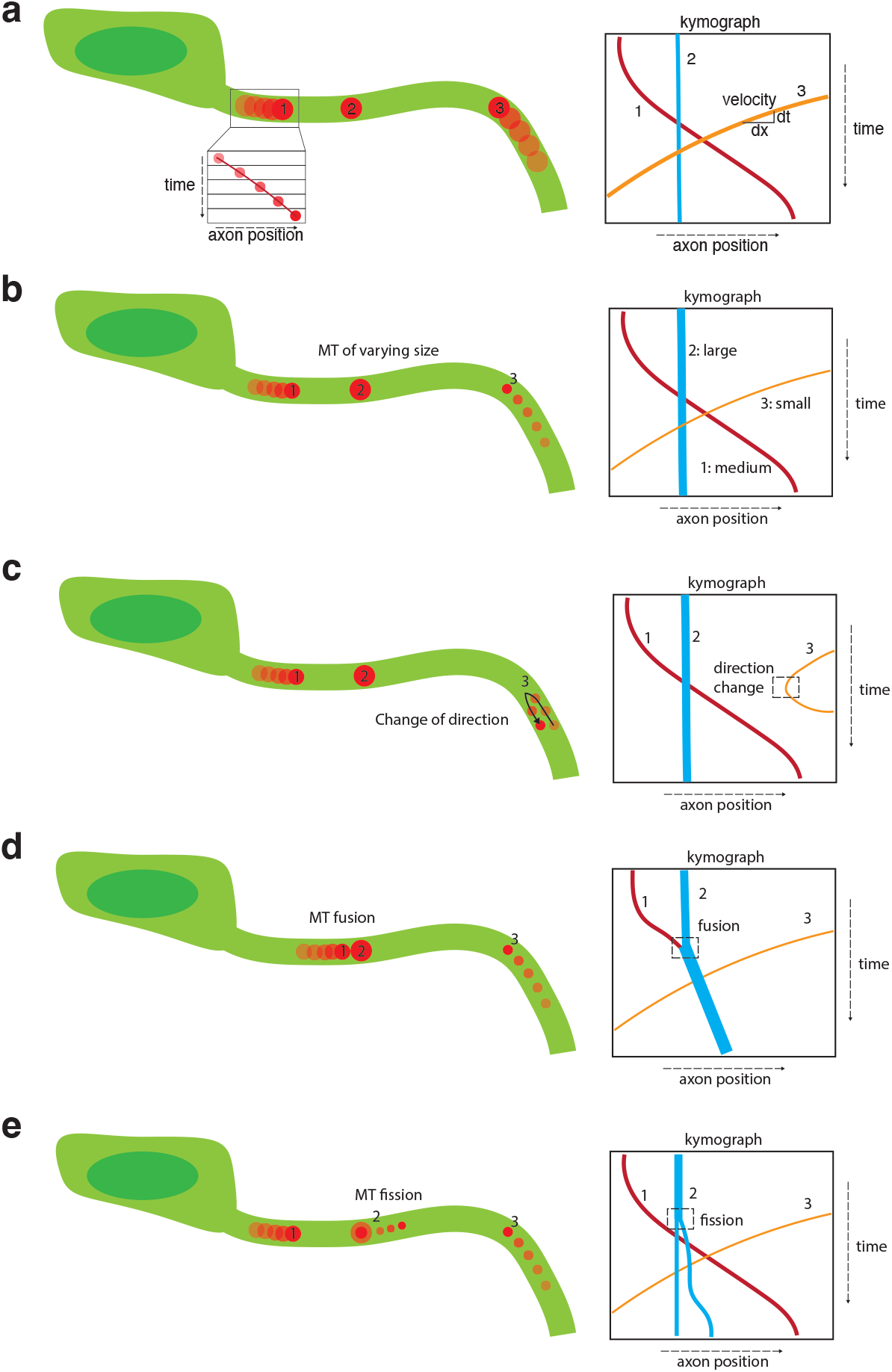
Schematic diagram of mitochondrial dynamics captured in kymographs. (A) The instantaneous velocity of each mitochondrion is captured through the slope of its corresponding line. (B) Mitochondrial size is captured in kymographs through line thickness. (C) Mitochondrial changes in direction appear as local minima or maxima within a given track. (D) Mitochondrial fusion events appear as two lines merging into one going from top to bottom. (E) Mitochondrial fission events appear as one line splitting into two going from top to bottom.

## Supplementary Information

Videos are provided as separate files and are not embedded in this PDF.

**Supplementary Video 1** MFN2 protein structure with positions of four amino acid substitutions labeled. File: `videos/mfn2_structure`.

**Extended Data Fig. 2.**
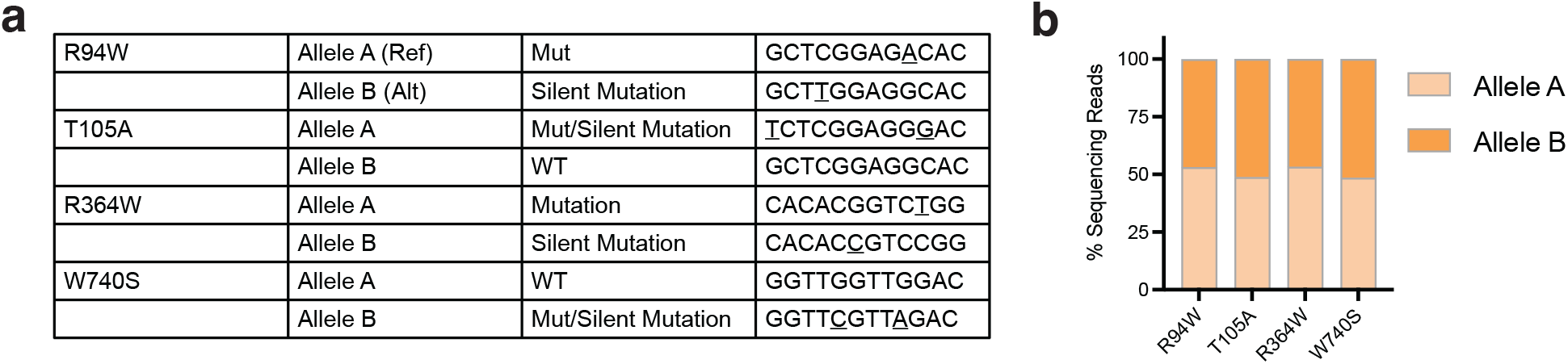
Sequence validation of all cell lines. (A) Consensus sequence for each allele at target site. (B) Allelic fraction quantified from Next Generation Sequencing reads.

**Supplementary Video 2** Representative example of an R364W axon segment that is enriched for C0 DINOv3 patches. Axons are green and mitochondria are red. The title for each video denotes the experiment ID, well ID, true genotype, kymograph ID, axon segment length, the predicted R364W probability from the KymoClass instance classifier (Extended Data Fig. 7), and the frequency of each DINOv3 patch cluster. Tick labels are noted in units of *µ*m. The time is denoted in both frame number and seconds at each frame of the video. File: `videos/D11_F03_kymo_47_len_176.mp4`.

**Supplementary Video 3** Representative example of a WT axon segment that is enriched for C1 DINOv3 patches. Axons are green and mitochondria are red. The title for each video denotes the experiment ID, well ID, true genotype, kymograph ID, axon segment length, the predicted R364W probability from the KymoClass instance classifier (Extended Data Fig. 7), and the frequency of each DINOv3 patch cluster. Tick labels are noted in units of *µ*m. The time is denoted in both frame number and seconds at each frame of the video. File: `videos/D11_B11_kymo_52_len_104.mp4`.

**Extended Data Fig. 3.**
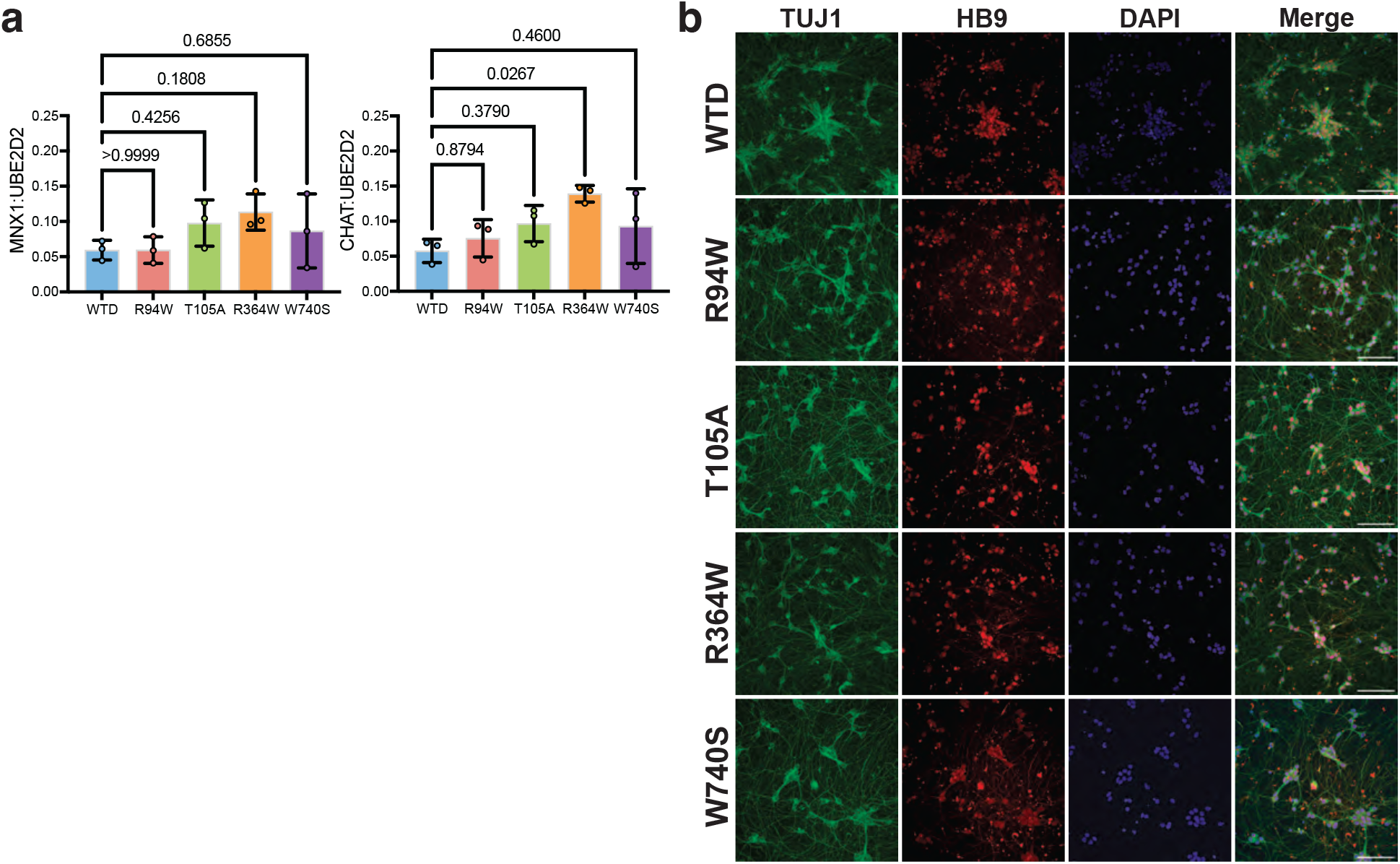
Differentiation state of all cell lines. (A) Digital PCR gene expression assay for motor neuron-specific transcription factor, *MNX1*, and the mature motor neuron marker, *CHAT*, normalized to *UBE2D2* expression. (B) Immunofluorescent-stained i^3^ motor neurons. Scale bar = 20 *µ*m. P-values were calculated by a one-way ANOVA, followed by Dunnett’s multiple comparison test against WT.

**Extended Data Fig. 4.**
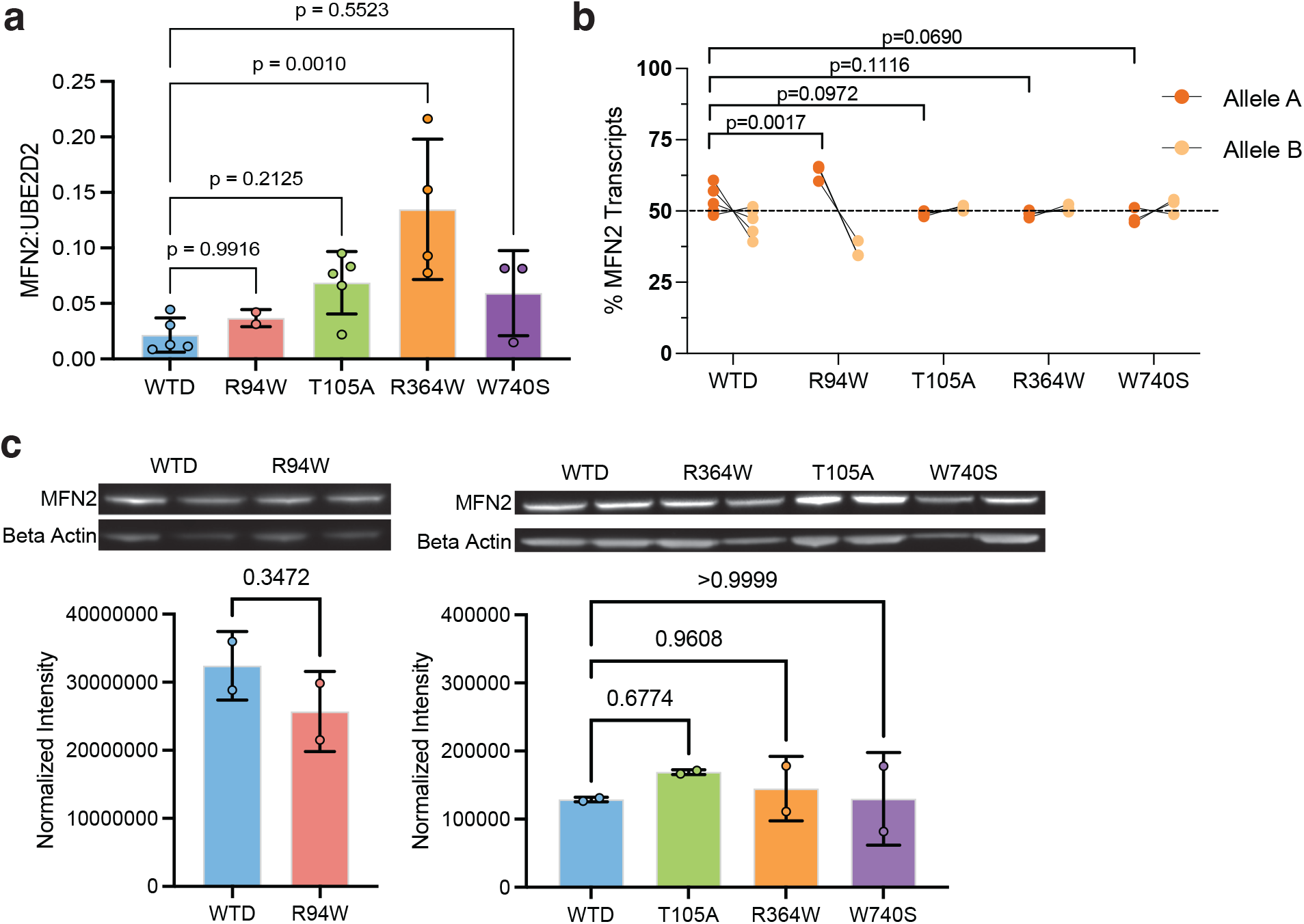
MFN2 expression of all cell lines. (A) Digital PCR gene expression assay for total *MFN2* normalized to *UBE2D2* expression. (B) Digital PCR assay for relative expression of *MFN2* transcripts. (C) Total MFN2 protein expression measured by western blot, normalized to *β* -actin. P-values were calculated by a one-way ANOVA, followed by Dunnett’s multiple comparison test against WT.

**Extended Data Fig. 5.**
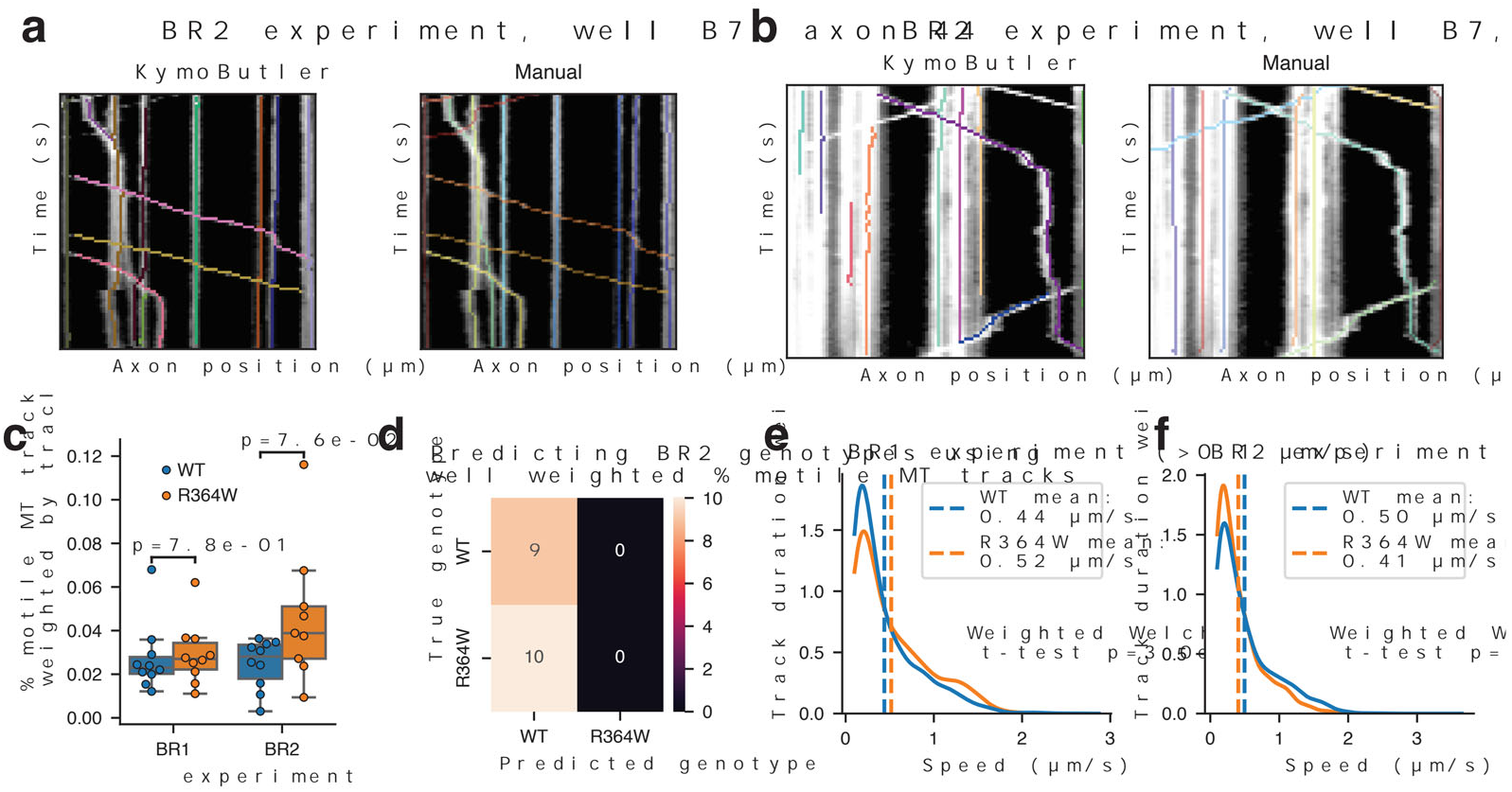
Representative comparisons between KymoButler and manual track segmentations and adjusting for KymoButler track duration. (A-B) Representative examples of KymoButler versus manual track segmentations overlaid onto raw kymographs. (C) Percentage of motile mitochondria tracks per well after adjusting for track duration, split by genotype and experiment. A two-sided t-test was performed to compare the two genotypes within each experiment. (D) Confusion matrix of predicting the genotype of BR2 wells using a logistic regression classifier trained on the track-duration-weighted percent motile values from BR1 wells. (E-F) Track-duration-weighted kernel density estimation of mitochondrial speed for each genotype in the (E) BR1 and (F) BR2 experiments. Weighted Welch’s t-tests were performed to compare the two genotypes within each experiment.

**Extended Data Fig. 6.**
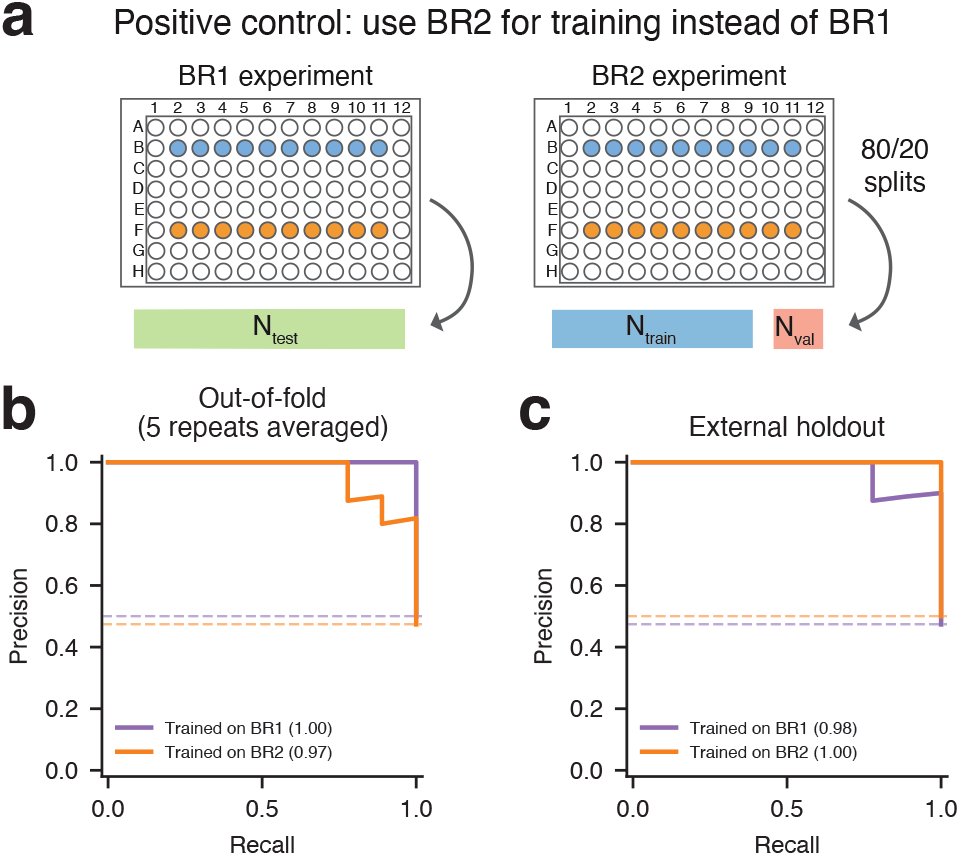
Positive control plate-swap experiment. (A) Instead of using BR1 wells for training and validation and BR2 wells as the held-out test set, we trained a separate KymoClass model that used BR2 wells for training and validation and BR1 wells as the held-out test set. (B-C) Precision-recall curves for predicting genotype on (B) the out-of-fold wells in the biological replicate used for training and (C) the wells in the holdout test set. Curves are shown for KymoClass models trained on BR1 and BR2. The AUC of each model is shown in parentheses.

**Extended Data Fig. 7.**
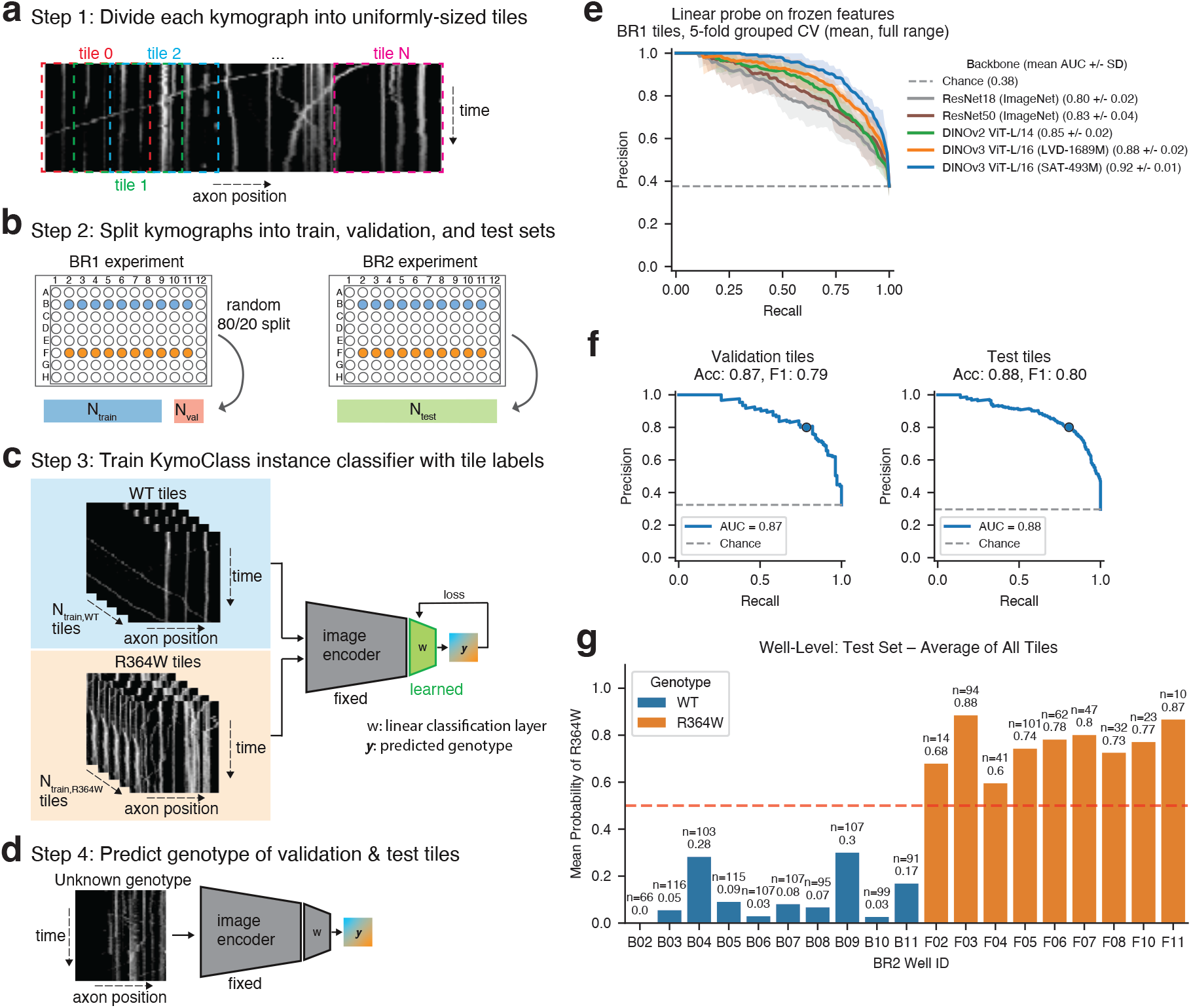
A KymoClass instance classifier predicts the genotype of kymograph tiles. (A-D) Overview of model training and evaluation scheme when predicting the genotype of individual tiles. (A) Each kymograph is divided into uniformly sized, overlapping tiles. Each tile contains all kymograph time points but not all positions of the axon. (B) Tiles from the BR1 experiment are randomly split into train and validation sets using an 80/20 split, while all tiles from the BR2 experiment are part of the test set. (C) The KymoClass instance classifier model takes kymograph tiles as input, passes them into a frozen image encoder, and trains a linear classification head to predict the genotype label of the tile. (D) To evaluate the model’s accuracy, we pass validation and test set tiles into the trained classifier, comparing the predicted R364W probability with the true genotype label. (E) Precision-recall curves across grouped 5-fold cross validation of BR1 tiles for five image encoder backbones. Each backbone’s across-fold mean curve is a solid line and the full range across folds is a shaded band. AUC means *±* standard deviations across folds are shown in parentheses. The DINOv3 SAT-493M model is selected for all downstream analysis. (F) Precision-recall curves for predicting tile genotype in the BR1 validation set (left) and BR2 test set (right). The accuracy and R364W-class F1 scores in subplot titles represent performance at the 0.5 probability cut, denoted by the single point along each curve. (G) Mean predicted R364W probability across all tiles of each BR2 test well, with bars colored by true genotype. Each bar is annotated with its mean value and the number of tiles in that well.

**Extended Data Fig. 8.**
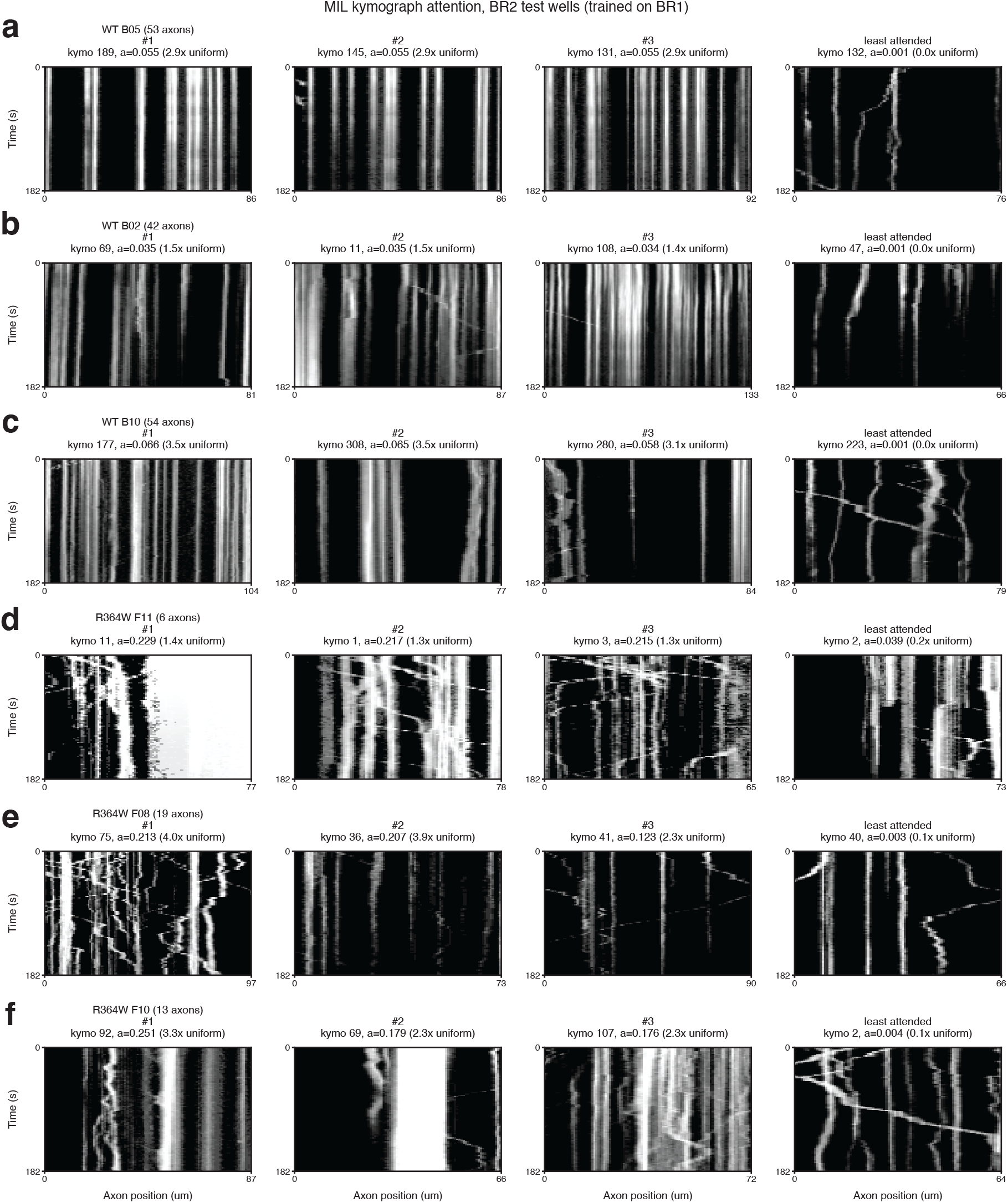
Kymographs with high and low attention scores in the WT versus R364W KymoClass MIL model. (A) The three kymographs with the highest attention scores (left) and one kymograph with the lowest attention score for WT well B05. The raw attention score and attention relative to the uniform weight are noted in the title of each kymograph. X-axis ticks represent the length of the kymograph in units of *µ*m. Kymographs are from the BR2 test set using a classifier trained on BR1. (B-C) Same as above except for WT wells (B) B02 and (C) B10. (D-F) Same as above except for R364W wells (D) F11, (E) F08, and (F) F10.

**Extended Data Fig. 9.**
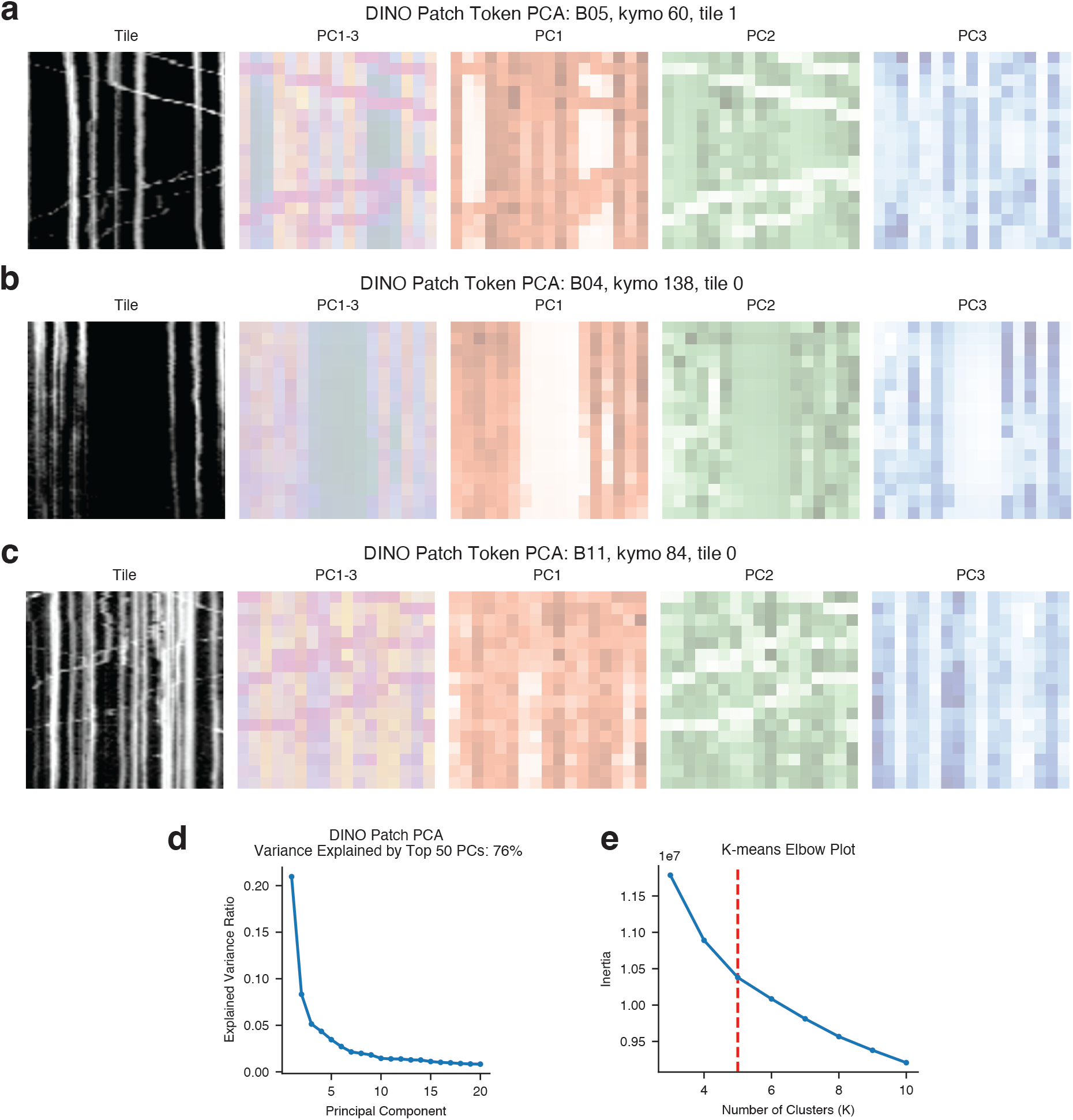
DINOv3 patch token embeddings can be used to consistently map all parts of a kymograph without supervision. (A-C) Results from running PCA on an *N ×* 1024 matrix where *N* represents all patches across all kymographs. Three representative kymograph tiles are shown. The raw kymograph is shown in black and white on the left. All 3 PCs are shown together in one image using PC1 scores as the red channel, PC2 scores as the green channel, and PC3 scores as the blue channel. The three rightmost columns show one PC at a time. (D) Variance explained by the top 20 PCs when running PCA on the DINOv3 patch embeddings with up to 50 PCs. The cumulative variance explained across all 50 PCs is shown above. (E) K-means elbow plot that shows the inertia of clustering the DINOv3 patch embeddings with different numbers of K-means clusters *K*. A red dashed line is shown at *K* = 5 as this is used for subsequent analysis.

**Extended Data Fig. 10.**
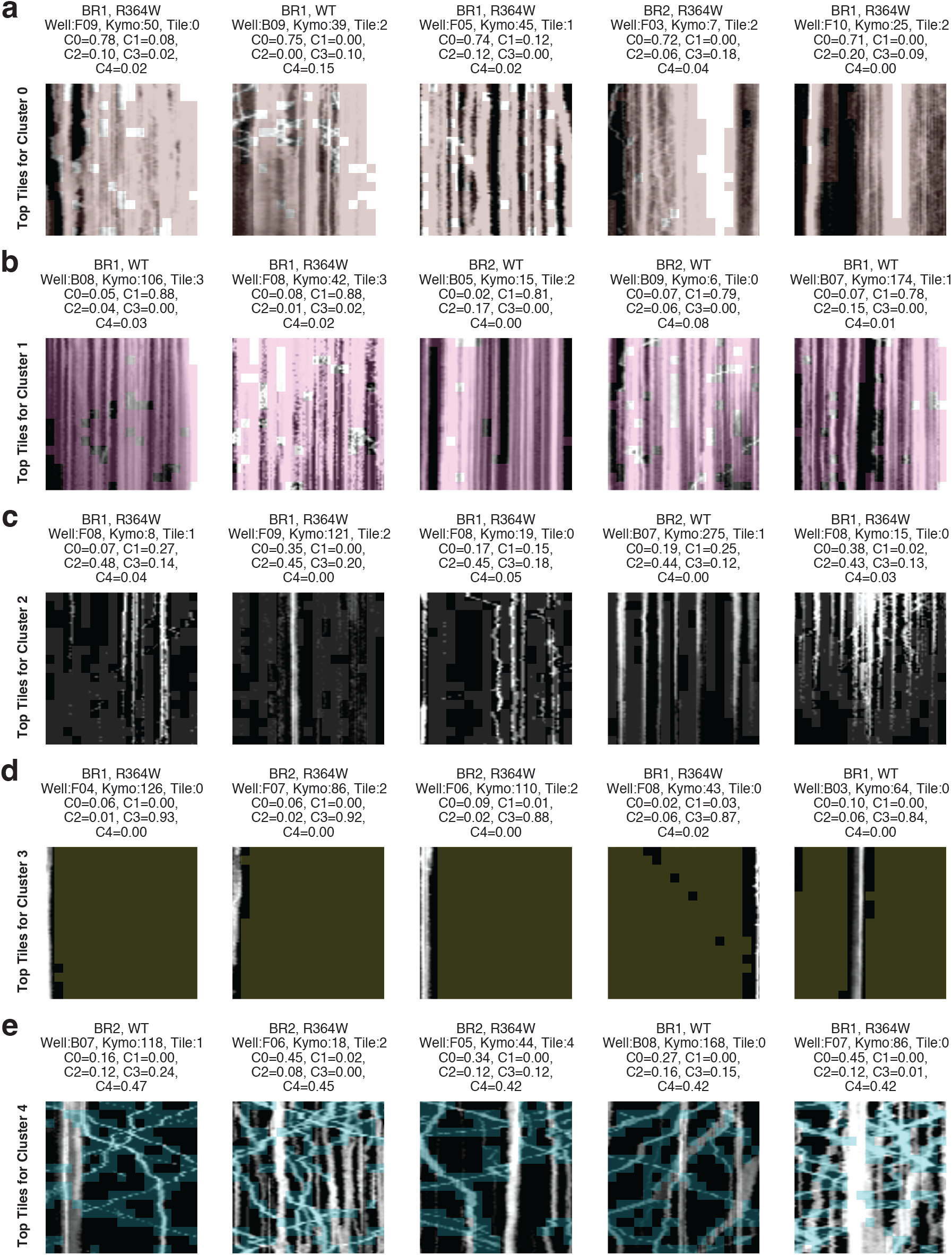
Kymograph tiles with the highest frequency of each DINOv3 patch cluster. (A) The five tiles with highest frequency of patch cluster C0. Each kymograph is shown in black and white, with patches belonging to C0 being superimposed with blue coloring. Patches with no superimposed color belong to a cluster other than C0. Subplot titles show the frequency of each patch token embedding cluster within a tile. At most one tile per kymograph is plotted. (B-E) Same as panel A except for clusters C1–C4, respectively.

**Extended Data Fig. 11.**
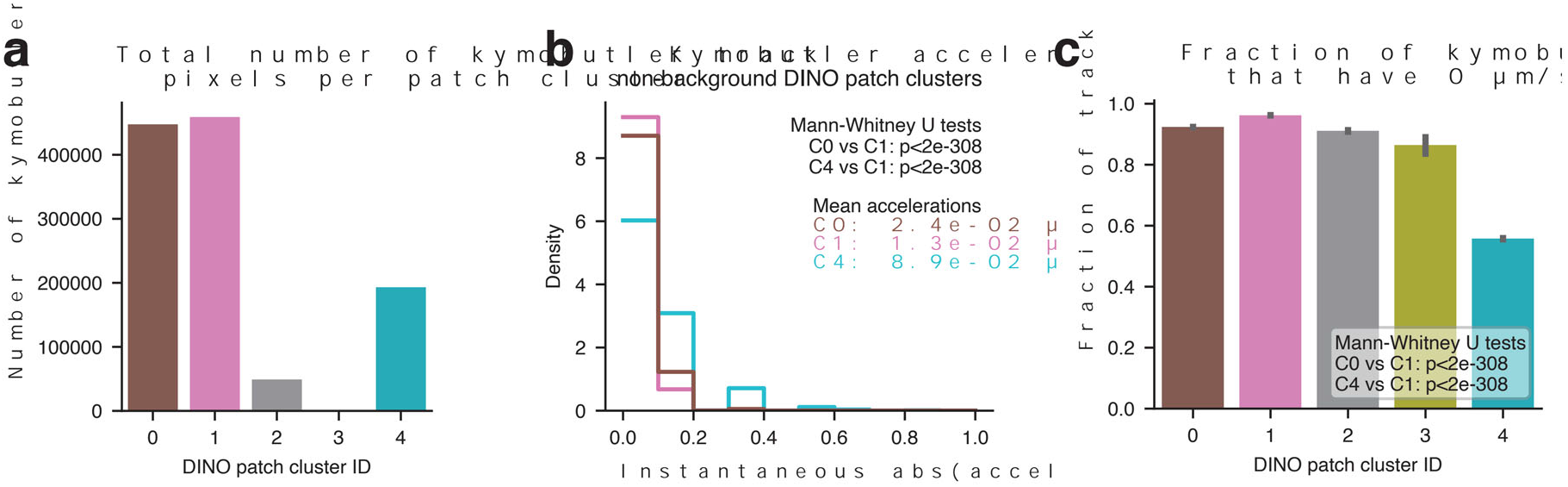
Additional summary statistics when using KymoButler tracks to interpret DINOv3 patch clusters. (A) Total number of KymoButler track pixels that appear within each DINOv3 patch cluster. (B) Distribution of instantaneous absolute acceleration for the three non-background DINOv3 patch clusters. (C) Fraction of track pixels with instantaneous velocity of 0 *µ*m/s within each DINOv3 patch cluster. Error bars represent 95% confidence intervals across 1000 bootstrap samples. Mann-Whitney U tests are computed for the other non-background clusters (C0 and C4) relative to C1.

**Extended Data Fig. 12.**
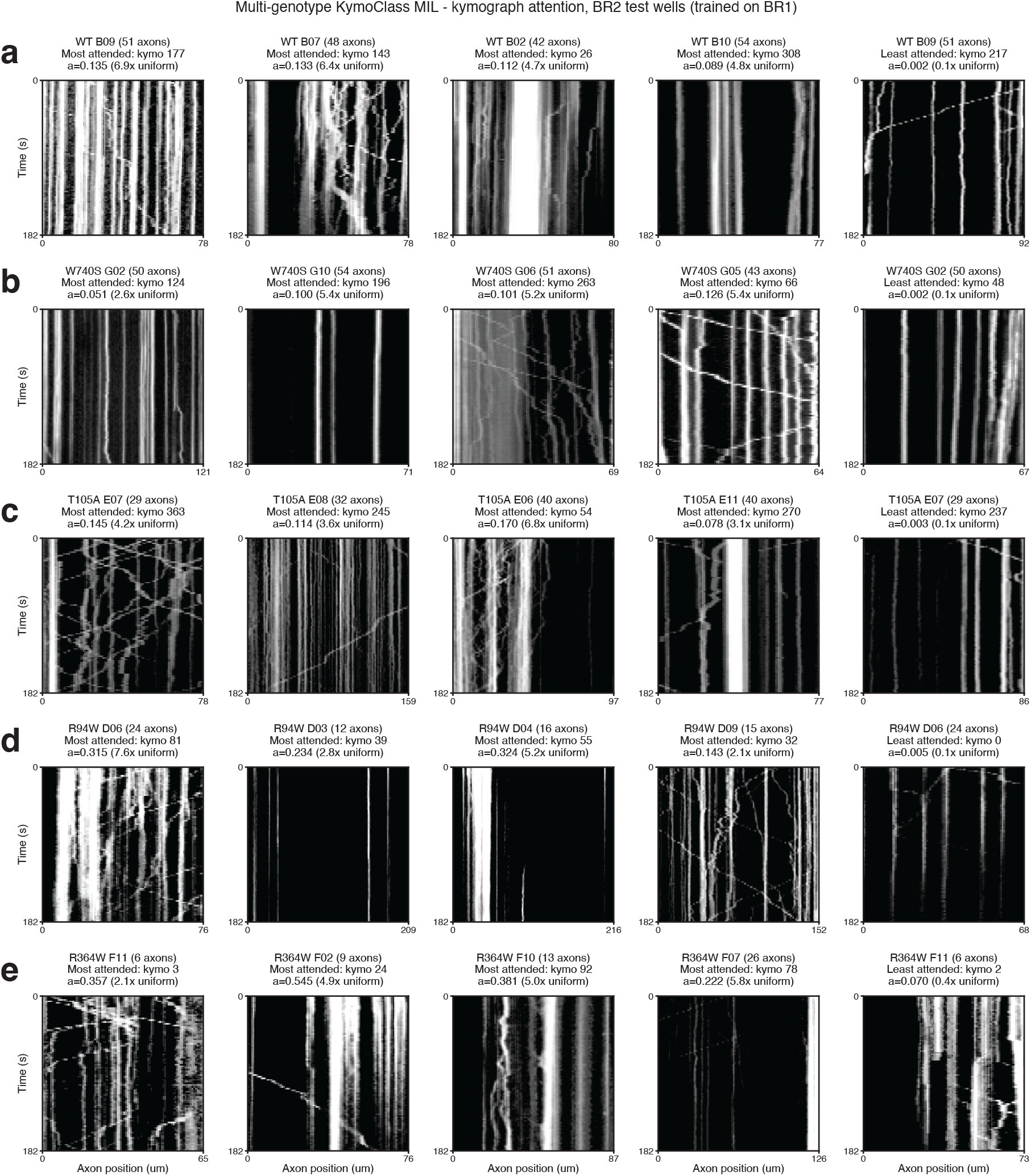
Kymographs with high and low attention scores in the multi-genotype KymoClass MIL model. (A) The most attended WT kymograph of each of four WT wells (left) and the least attended kymograph of one of those wells (right), according to the multi-genotype KymoClass model. The well ID, kymograph ID, raw attention score, and attention relative to the uniform weight are noted in the figure title. X-axis ticks represent the length of the kymograph in units of *µ*m. Kymographs are from the BR2 test set using a classifier trained on BR1. (B-E) Same as above except for genotypes (B) W740S, (C) T105A, (D) R94W, and (E) R364W.

**Extended Data Fig. 13.**
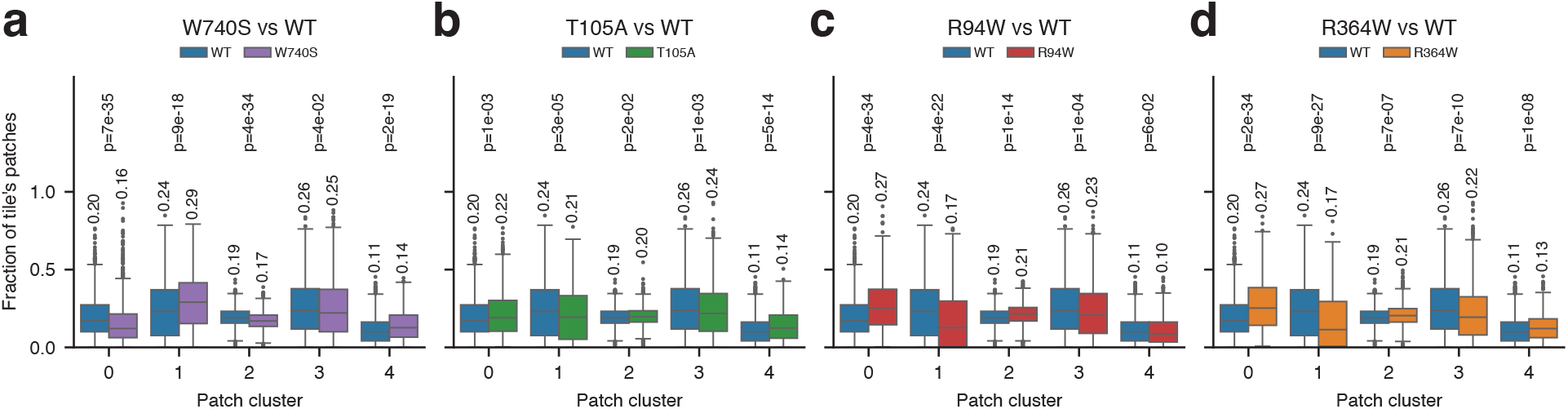
Comparing the abundance of each DINOv3 patch cluster between each. *MFN*2^mut^ **genotype and WT.** (A) Fraction of each tile’s patches belonging to a given DINOv3 patch cluster, compared between W740S tiles and WT tiles with both biological replicates pooled. Each box summarizes the tiles of one genotype for one cluster, and that box’s mean fraction is annotated above it. Each p-value is from a two-sided Mann-Whitney U test comparing the tiles of W740S wells against the tiles of WT wells, after Benjamini-Hochberg false discovery rate correction across the 20 tests shown in this figure. (B-D) Same as panel A, for the (B) T105A, (C) R94W, and (D) R364W mutations.

